# Haplotype phased genome of ‘Fairchild’ mandarin highlights influence of local chromatin state on gene expression

**DOI:** 10.1101/2024.01.20.575729

**Authors:** Isaac A. Diaz, Talieh Ostovar, Jinfeng Chen, Sarah Saddoris, Robert J. Schmitz, Susan R. Wessler, Jason Stajich, Danelle K. Seymour

**Affiliations:** Department of Botany and Plant Sciences, UC Riverside; Department of Microbiology and Plant Pathology, UC Riverside; Department of Genetics, University of Georgia

## Abstract

**Background:** Cis-regulatory sequences control gene expression through the coordinated action of transcription factors and their associated partners. Both genetic and epigenetic perturbation of cis-regulatory sequences can lead to novel patterns of gene expression. Phased genome assemblies now enable the local dissection of linkages between cis-regulatory sequences, including their epigenetic state, and gene expression to further characterize gene regulation in heterozygous genomes.

**Results:** We assembled a locally phased genome for a mandarin hybrid named ‘Fairchild’ to explore the molecular signatures of allele-specific gene expression. With genome phasing, genes with allele-specific expression were paired with haplotype-specific chromatin states, including levels of chromatin accessibility, histone modifications, and DNA methylation. We found that 30% of variation in allele-specific expression could be attributed to haplotype associated factors, with allelic levels of chromatin accessibility and three histone modifications in gene bodies having the most influence. Structural variants in promoter regions were also associated with allele-specific expression, including specific enrichments of hAT and MULE-MuDR DNA transposon sequences. Mining of cis-regulatory sequences underlying regions with allelic variation in chromatin accessibility revealed a paternally-associated sequence motif bound by ERF48, a target of the Polycomb repressive complex 2 (PRC2), and sequence similarity of this motif corresponded to local levels of H3K27me3, a signature of PRC2 activity.

**Conclusions:** Using a locally phased assembly of a heterozygous citrus cultivar, we dissected the interplay between genetic variants and molecular phenotypes with the goal of revealing functional cis-regulatory sequences and exploring the evolution of gene regulation.

## Introduction

Spatiotemporal modulation of gene expression programs is essential for cellular function, organismal development, and environmental responsiveness. Specific gene expression patterns are achieved through control of cis-regulatory modules (CRMs), which include sequences that influence the timing, magnitude, and frequency of transcription through the coordinated action of transcription factors (TF) and other binding partners. Local chromatin environment influences the ability of TFs to bind CRMs and promote or repress gene expression and variation in chromatin state at CRMs underlies variation in gene expression, a major source of phenotypic novelty [1].

Interspecific hybridization has the potential to generate unique combinations of cis-regulatory activity that result in novel phenotypes [2]. Allelic cis-regulatory variation may be due to either genetic perturbation of TF binding motifs in CRMs or through differential accessibility of CRMs achieved by epigenetic modulation of chromatin state. In both cases, transcription will be imbalanced between the two alleles as a result of differential regulation of CRMs. The comparison of local chromatin state and gene expression between first generation hybrids and their parents can reveal the relative contribution of cis-acting variation, including genetic and epigenetic variation in CRMs, and trans-acting variation caused by factors encoded in other genomic locations. For example, chromatin packing in an interspecific hybrids between *Arabidopsis thaliana* and *Arabidopsis lyrata* revealed that chromatin packaging was mostly unchanged for the *A. lyrata* genome, but varied substantially for the *A. thaliana* genome, with increased chromatin compaction of *A. thaliana* chromosomes in the interspecific hybrid [3]. Similarly, more *A. thaliana* genes were differentially regulated in the interspecific hybrid than *A. lyrata* genes although the total number of affected genes was relatively small [3].

In the genus *Citrus*, the hybridization of three ancestral species, citron (*C. medica*), pummelo (*C. maxima*), and mandarin (*C. reticulata*), has produced many hybrids with novel phenotypes including sweet oranges (*C. sinensis*), lemons (*C. limon*), and grapefruits (*C. grandis*) [4]. There are several examples where novel phenotypes in cultivated citrus hybrids are the result of cis-regulatory variation [5,6]. For example, in blood oranges the insertion of a Copia-like retrotransposon upstream of the *Ruby* gene introduces one or more CRMs that modulate gene expression in response to temperature [5]. Upregulation of *Ruby* transcripts in response to cold temperatures results in anthocyanin accumulation in the fruit [5]. Apomixis, another commercially important trait, is also associated with cis-acting regulatory variation introduced by a transposon sequence [6,7]. The production of apomictic seed, or seed genetically identical to the mother plant, is controlled in part by a gene encoding a RWP-RK domain–containing protein (*CitRWP*) with the insertion of a miniature inverted repeat transposable element (MITE) in its promoter linked to enhanced gene expression in *Citrus* and its relative *Fortunella hindsii* [6]. These are only two examples where structural variants alter gene expression patterns by perturbing CRMs, including through the introduction of a new CRM to regulate the cold-responsiveness of *Ruby*. The impact of structural variation on gene expression is likely much higher and an extensive inventory of structural variation in tomato revealed that 7.3% of genes with structural variants (SVs) in cis-regulatory regions were differentially expressed [8]. There is accumulating evidence that perennial, clonally propagated crop species, like *Citrus,* harbor extensive structural variation between haplotypes [6,9], but the impact of these variants on phenotypic variation, including potential effects on cis-regulatory activity, remains to be characterized at the genome-wide level.

CRMs include sequences with the potential to enhance and repress expression of neighboring genes. In plants, CRMs can be broadly categorized as core promoters, enhancers, or silencers. Core promoters contain the minimal sequence required for transcription initiation, while enhancers and silencers contain elements that can be bound by transcription factors (TFs) to increase or decrease gene expression, respectively. Core promoters are usually within 50-100 bp of the transcription start-site while enhancers and silencers can be located much farther from the gene targeted for regulation [10]. For example, the maize enhancer *Hopscotch* is located about 70 Kb from its target *teosinte branched 1* [11]. Core promoters, enhancers, and silencers may exhibit distinct patterns of chromatin accessibility and histone modifications depending on whether they are in a poised, active, or inactive state [10]. Active promoters are characterized by chromatin accessibility and activating histone modifications such as H3K4me3 and histone acetylation [12–16]. In contrast, inactive promoters have reduced chromatin accessibility and can be marked by H3K27me3, a repressive histone modification catalyzed by Polycomb repressive complex 2 [17]. One challenge for studying cis-regulatory variants is the identification of active regulatory regions from the massive amount of non-coding DNA in the genome. The genome-wide identification of accessible chromatin regions (ACRs), or regions of the genome that are not tightly packaged around nucleosomes, using ATAC-seq [18] narrows the portion of intergenic space that contains likely CRMs. A survey of angiosperms revealed that only 0.22-6.5% of the genome resides in accessible chromatin regions in leaf tissue [19]. The chromatin landscape of ACRs has been used to infer their role in activating or repressing transcription of neighboring genes [15,19].

Here, we investigate the contribution of cis-regulatory variation to generating novel patterns of gene expression in a cultivated mandarin hybrid named ‘Fairchild’. We quantify allelic variation in both chromatin state and gene expression in this hybrid. To directly link local chromatin variation to gene expression we generated a locally phased reference genome of ‘Fairchild’. Genes with imbalanced allelic expression served as a focal point to explore the contribution of the chromatin and epigenetic landscape to gene expression variation. We found that allele-specific expression can be predicted, in part, by local chromatin state, with chromatin accessibility and histone modifications of gene bodies being the most influential factors. Local structural variation was also linked to allele-specific variation in gene expression, with hAT and MULE-MuDR DNA transposons enriched at genes with divergent expression between haplotypes. Finally, we identified differential ACRs between haplotypes, and mined these sequences to identify candidate motifs underlying differential gene expression patterns. We show that cis-regulatory variation between haplotypes is reflected in the chromatin landscape and that TE-associated genetic variants may underlie expression divergence in this cultivated citrus hybrid.

## Results

### The landscape of accessible chromatin in a haplotype-phased genome

We sought to investigate the chromatin dynamics at cis-regulatory sequences and the influence of those sequences on gene regulation in the heterozygous genome of ‘Fairchild’. To address this we first *de novo* assembled the genome of ‘Fairchild’ using 69 Gb of PacBio long-read sequences (Materials and methods). The *de novo* assembly was scaffolded using a *Bionano* optical map and anchored to chromosomes based on existing genetic maps (Materials and methods). The resulting ‘Fairchild’ genome assembly is 365 Mb in length (N50 = 42.1 Mb), with 95% of the genome contained in 9 chromosomes (Additional File 2: Fig. S1). Cis-regulatory modules (CRMs) typically include an array of transcription factor (TF) binding sites, with the majority of TFs only binding to DNA sequence in open chromatin conformations [20–22]. Putative CRMs in ‘Fairchild’ were identified by mapping accessible chromatin regions (ACRs) using ATAC-seq of nuclei extracted from leaf tissue. A total of 9,172 ACRs spanning 7.3 Mb (FRiP = 0.13) were identified (Materials and methods, Additional File 1: Tables S1, S2). The sequences underlying these ACRs span 2% of the ‘Fairchild’ genome, similar to other plant species [19].

Recent work has demonstrated the importance of distal cis-regulatory sequences in controlling gene regulation in plants [15]. In maize, for example, 32% of ACRs are located more than 2 Kb, and often more than 20 Kb, from their nearest gene [19]. Based on this, ‘Fairchild’ ACRs were classified as genic, proximal, or distal based on proximity to their nearest gene (overlapping, within 2 Kb, or further than 2 Kb, respectively). The majority (53.6%) of ACRs are genic, while 33.7% of ACRs are proximal, and 17.3% are distal to their nearest gene (Additional File 1: Table S2). We then determined the genome-wide distribution of three activating histone modifications (H3K4me3, H3K36me3, H3K56ac) and one repressive modifications (H3K27me3), using ChIP-seq (Materials and methods) (Additional File 1: Table S3-S6). Unsupervised clustering of histone modifications at ACRs revealed similar patterns observed in other plant species [19] (Additional File 2: Fig. S2, S3,S4). When focusing on only distal ACRs (> 2 Kb from nearest gene), a distinct cluster of ACRs enriched for the repressive modification H3K27me3 emerges that is not present for genic and proximal ACRs. (Additional File 2: Fig. S3) It is possible that these distal ACRs marked by repressive modifications represent poised enhancers [23,24], although not much is known about histone modification of poised enhancers in plants.

To define the extent of local and distal cis-regulatory variation, we generated locally phased haplotypes with 10x Genomics Linked-Reads. In total, 1.53 phased variants, including small indels (< 50 bp) were used to define 1,015 phase blocks (N50 = 1.57 Mb). Phase blocks spanned 88.04% of the assembled genome length and included 91.2% of annotated genes, with 45.4% of haplotype blocks including one or more genes (Fig. 1a). In other plant species, distal CRM have been identified more than 60 Kb from their target gene [11], but there are typically no intervening coding gene sequences [15]. This suggests that allele-specific chromatin dynamics at both local and distal CRM can be assayed for 91.2% of gene sequences in ‘Fairchild’ based on the multi-megabase scale of haplotype phasing. SNP-based read phasing enabled the quantification of allelic gene expression, chromatin accessibility, histone modification, and DNA methylation (Fig. 1b). Allelic differences in whole-genome sequencing read mapping served as a control for read mapping bias (Materials and methods). This approach enables comparison of haplotype-specific signatures of chromatin landscape and gene expression in this hybrid genome.

**Fig 1:**
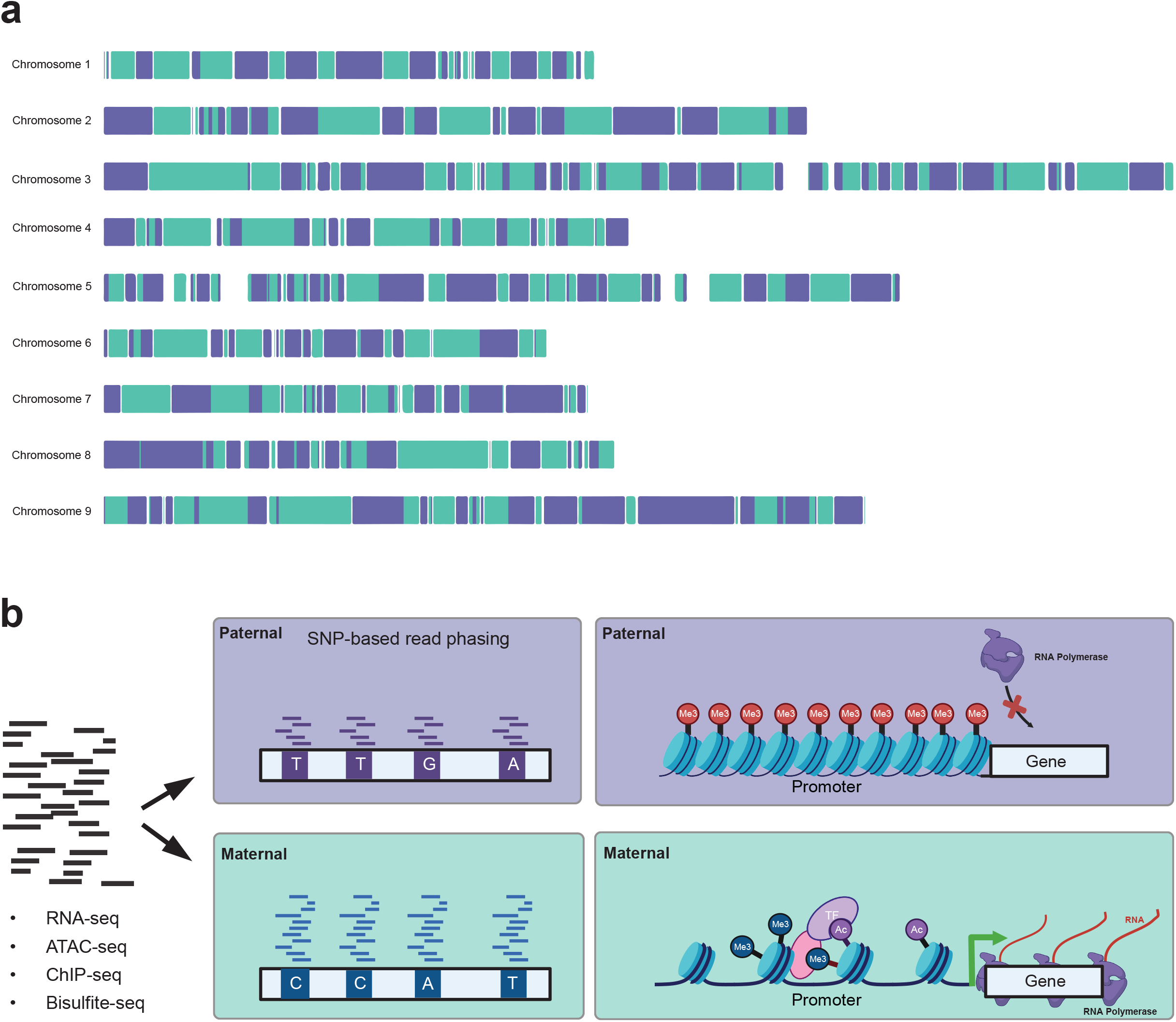
Framework for developing haplotype-resolved profiles of gene expression, chromatin accessibility, and epigenetic modifications**. a)** Phased haplotype blocks along each chromosome of the ‘Fairchild’ reference genome constructed using 10x Genomics Linked-Reads (n = 1,015, N50 = 1.57 Mb). Alternating colors indicate the distinct phase blocks. **b)** Diagram illustrating the strategy for SNP-based phasing of RNA-seq, ATAC-seq, ChIP-seq, and Bisulfite-seq reads into maternal and paternal haplotypes. **c)** Example of repressive (paternal) and active (maternal) chromatin landscapes in proximity to a gene with allele-specific expression (ASE).

To evaluate the extent of haplotype-specific control of gene expression we utilized phased SNPs to quantify allele-specific expression (ASE) by RNA-seq (Materials and methods). Overall, 23,407 genes are expressed in the leaf and 7,964 (34.02%) of those genes have sufficient levels of polymorphism (2 or more SNPs) for allele-specific analysis. Genes evaluated for allele specific expression contained 3 SNPs per 1,000 bp, on average. In total 2,071 genes with significant ASE consistent across two replicates were identified (Benjamini Hochberg FDR < 0.05) (Additional File 1: Tables S6, S7, S8). Genes with ASE were enriched for gene ontology terms related to metabolism and other enzymatic processes (Additional File 1: Table S9). Although genetic variation limits our ability to assay allele-specific patterns of expression across all expressed genes, ASE appears to be common in this subset, affecting 26% of genes with sufficient polymorphism and levels of expression. This set of ASE genes serve as the basis for investigating the dynamics of haplotype-specific cis-regulatory variation.

### Haplotype-specific chromatin dynamics correlate with allele-specific expression

Regulatory variants have the potential to cause haplotype-specific alterations in chromatin state through differential transcription factor and chromatin remodeling activity. We assigned ancestry to phased SNPs to represent allele-specific gene expression and chromatin state by parent of origin. ‘Fairchild’ was derived from a cross between a clementine mandarin and ‘Orlando’ tangelo and re-sequencing data from a clementine and ‘Orlando’ were used to assign ancestry to each phased SNP (Additional File 1: Table S10). Local phasing of haplotypes with ASE genes enabled us to examine the extent to which haplotype-specific variation in the chromatin landscape is associated with the expression of linked genes.

To address this, we assayed the landscape of four histone modifications (H3K4me3, H3K36me3, H3K56ac, H3K27me3), chromatin accessibility, and DNA methylation, including their haplotype-specific patterns, to dissect the relative contribution of each feature to gene expression. We then partitioned all seven metrics of chromatin status into four regions flanking each expressed gene: genic (including exon and introns), promoter (< 1 Kb from the transcription start site), upstream putative regulatory region (ACR within 5 Kb upstream of promoter), and downstream putative regulatory region (ACR within 5 Kb downstream of gene). For upstream and downstream regions, histone modifications and chromatin accessibility data were only incorporated if a putative regulatory region (ACR) was detected in the window of interest.

Focusing on the 2,071 genes with significant ASE, gene expression in transcripts per million (TPM) was then modeled using elastic-net regression with 56 factors (Additional File 1: Table S11), including each chromatin feature across the four regions. In addition to overall levels of histone modification and chromatin accessibility at each gene, haplotype-specific differences for each feature were included as factors. Additional factors included the presence of a SV in the promoter and the ratio of nonsynonymous and synonymous substitutions. After excluding genes that were outliers for whole-genome sequencing coverage (n = 162), five-fold cross validation was used to select the optimal parameters for elastic-net regression (Additional File 1: Table S12). 40% of the variation in overall gene expression was explained with 21 of the 56 factors (Fig. 2a; Additional File 1: Table S13, S14) with levels of genic chromatin accessibility (ATAC), genic H3K4me3, and genic H3K27me3 having the largest influence. Consistent with their roles in activating and repressing gene expression, H3K4me3 was positively correlated with overall expression, while H3K27me3 was negatively correlated with expression (Fig. 2a). When including all genes in the ‘Fairchild’ genome, 49 of the 56 factors explained 61.15% of variation in overall gene expression (Additional File 1: Table S15; Additional File 2: Fig. S5). The fact that almost all predictors in this model were determined to be significant suggests model overfitting. To address this, we selected a shrinkage parameter (lambda) that reduced model complexity while maintaining a prediction error within one standard error of the initial model (Additional File 1: Tables S15, S16; Additional File 2: Fig. S6). This reduced model included 15 of the 56 factors and explained 60.44% of the variance in overall gene expression. The factors with the greatest influence on gene expression of all genes are overall levels of genic H3K36me3, genic H3K56ac, genic H3K27me3, and chromatin accessibility of both promoters and genic regions (ATAC) (Additional File 1: Tables S16; Additional File 2: Fig. S7). In summary, genic levels of chromatin accessibility and H3K27me3 were 2 of the 5 top influential factors in models of overall expression for all genes (n= 30,724) and genes with ASE (n = 2,071), with the influence of open chromatin being more pronounced for genes with ASE.

**Fig 2:**
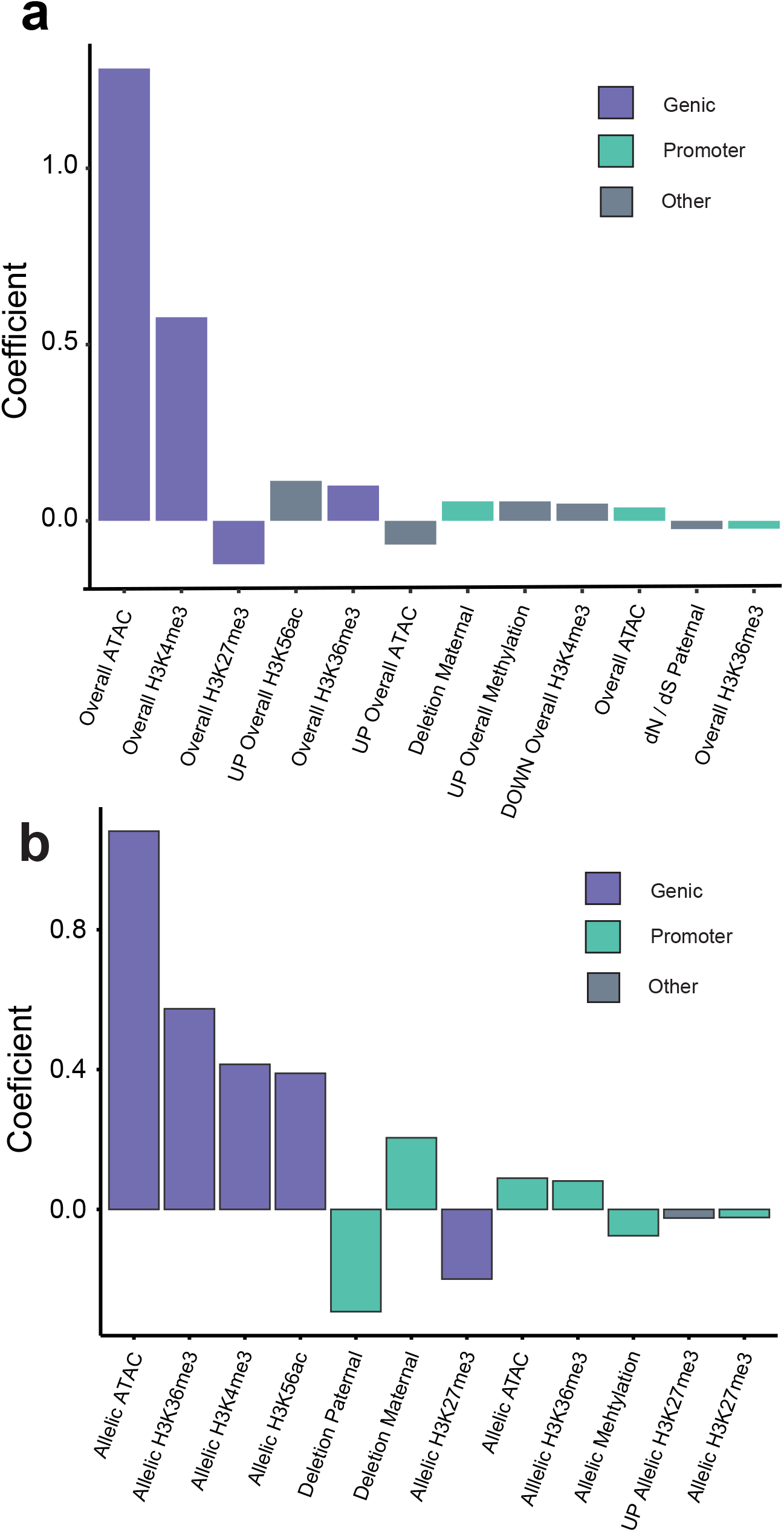
The relationship between gene-expression and chromatin features at genes with allele-specific expression. **a)** Coefficients of the top 12 factors for the model of overall gene expression (log_2_(TPM)) for 1,909 ASE genes ordered by magnitude (R = 0.40). Factors are partitioned by genomic region and colored to indicate whether they reside in genes, promoters (1 Kb of TSS), or upstream/downstream putative regulatory regions of the focal gene (ACRs present 5 Kb upstream of promoter / 5 Kb downstream of gene). **b)** Coefficients of the top 12 factors for the model of allelic gene expression (log_2_FC(Maternal_RNA_ / Paternal_RNA_) ordered by magnitude (R = 0.3069). Positive “allelic” factors indicate a positive association between maternal gene expression and maternal levels of a given data-type. “Overall” indicates that the predictor is the total level of a given data-type (ATAC, ChIP, Bisulfite). “Allelic” indicates that the predictor is the ratio of maternal:paternal levels of a given data-type (log_2_FC(Maternal / Paternal)).

A similar approach was used to quantify the contribution of each factor to allele-specific expression, or the log_2_ fold difference in the expression of the maternal allele (clementine) over the paternal allele (‘Orlando’). After removing ASE genes with apparent read mapping bias, 1,786 genes remained, with 30.69% of the variation in differential expression between alleles explained by 18 of the 56 factors (Fig. 2b; Additional File 1: Table S17). Allelic differences in levels of chromatin modifications were associated with ASE, unlike the model of overall expression for the set of 2,071 genes where overall levels of chromatin modifications were predictive of gene expression. Allelic chromatin accessibility, H3K36me3, H3K4me3, and H3K56ac in genic regions were the four strongest predictors of ASE and were all positively associated with ASE. The repressive modification H3K27me3 was inversely associated with allele-expression indicative of its repressive effect.

In addition to inferring the relative effect of chromatin features on allele-specific expression, we also examined the significance of their genomic location (genic, promoter, upstream putative regulatory region, downstream putative regulatory region) relative to each gene. Chromatin accessibility and histone modifications of genic regions have the strongest relationship with ASE, while promoter chromatin accessibility, promoter H3K36me3 and DNA methylation of promoters have more subtle effects (Fig. 2b; Additional file 1: Table S17). Allelic DNA methylation of promoters has an effect similar to that of genic H3K27me3, illustrating two modes of transcriptional repression. Additionally, the models reveal a significant effect of deletions in promoter sequences on gene expression, primarily in the model of ASE (Fig. 2b) but also in the model of overall expression (Fig. 2a). Using 10X Genomics Linked-Reads, we identified 2,481 deletions ranging from 41 bp to 11,084 bp (mean length = 332 bp) (Additional File 2: Fig S8). Although parentage can be assigned to each lesion, the designation as a ‘deletion’ is in reference to the ‘Fairchild’ consensus sequence and the ancestral state (i.e. inserted or deleted allele) is unknown. Here, a promoter deletion in one haplotype is associated with increased expression of the corresponding haplotype, with coefficients of similar size for paternal and maternal deletions (Fig. 2b). Since the model response is represented as the ratio of maternal vs paternal gene expression, the opposite effects of paternal and maternal deletions indicate their effect on the deletion containing allele (Fig. 2b). Overall the chromatin landscape in genic regions and promoters is more associated with ASE than distal regulatory regions. Together, these analyses demonstrate the extent to which allelic differences in histone modifications, chromatin accessibility, and DNA methylation underlie ASE.

### Promoter structural variation linked to differences in allele-specific expression

Genetic variants in gene promoter sequences may alter the binding affinity of regulatory factors, serving as a source of cis-regulatory variation. Since multiple functional elements reside in promoter regions, the effect of a single SNP may be small. In contrast, structural variants may alter the sequence or position of cis-regulatory modules and have potentially larger effects on gene expression. In ‘Fairchild’, the promoters of genes with ASE are significantly enriched for deletions (1,000 permutations, p < 0.001) (Additional File 2: Fig. S9). Additionally, promoter structural variants of genes with ASE are enriched for Type I DNA transposable elements (TEs), with the hAT and MULE-MuDR DNA transposons being the most abundant (Additional File 2: Figure S10). To further examine the relationship between SVs and ASE we identified 1,654 phased deletion:ASE gene pairs for which the gene and its nearest deletion are in the same phase block (Fig. 3a). Genes with a deletion in their promoter (n=104) had significantly higher expression of the deletion-containing allele (Hedges G effect-size 0.378) compared to ASE genes without promoter deletions (1,000 permutations, p < 0.001) (Fig. 3a). One example of this is the elevated expression of the maternal allele for a predicted aldehyde dehydrogenase gene (Additional File 1: Table S18). There is a 452 bp structural variant ∼900 bp upstream of the gene that corresponds with dramatically higher expression of the maternal allele (log_2_FC = 2.53) (Fig. 3b). Because the ancestral state of each structural variant is unknown, it could be that expression of the paternal allele, in this case, is associated with an insertion that reduces expression. These inserted sequences could disrupt cis-regulatory elements, or introduce repressive elements. A motif enrichment analysis of sequence removed in the ‘deleted’ allele of the 104 promoter:gene pairs identified four significantly enriched motifs, with the most abundant (n=28) matching the binding site of zinc-finger protein 1 (AZF1), a zinc-finger protein that acts as a transcriptional repressor [25] (Additional File 1: Table S19). For these deletion:gene pairs there was no significant effect of AZF1 motif presence on allele-specific expression, but the sample size was relatively small (10,000 permutations, p = 0.2717) (Additional File 2: Figure S11). However, 5 of the 10 ASE genes with the highest expression bias towards the deletion-containing allele included the canonical AZF1 motif (Fig. 3c). These observations suggest a possible role for repressive elements in structural variants linked to allele-specific expression.

**Fig 3:**
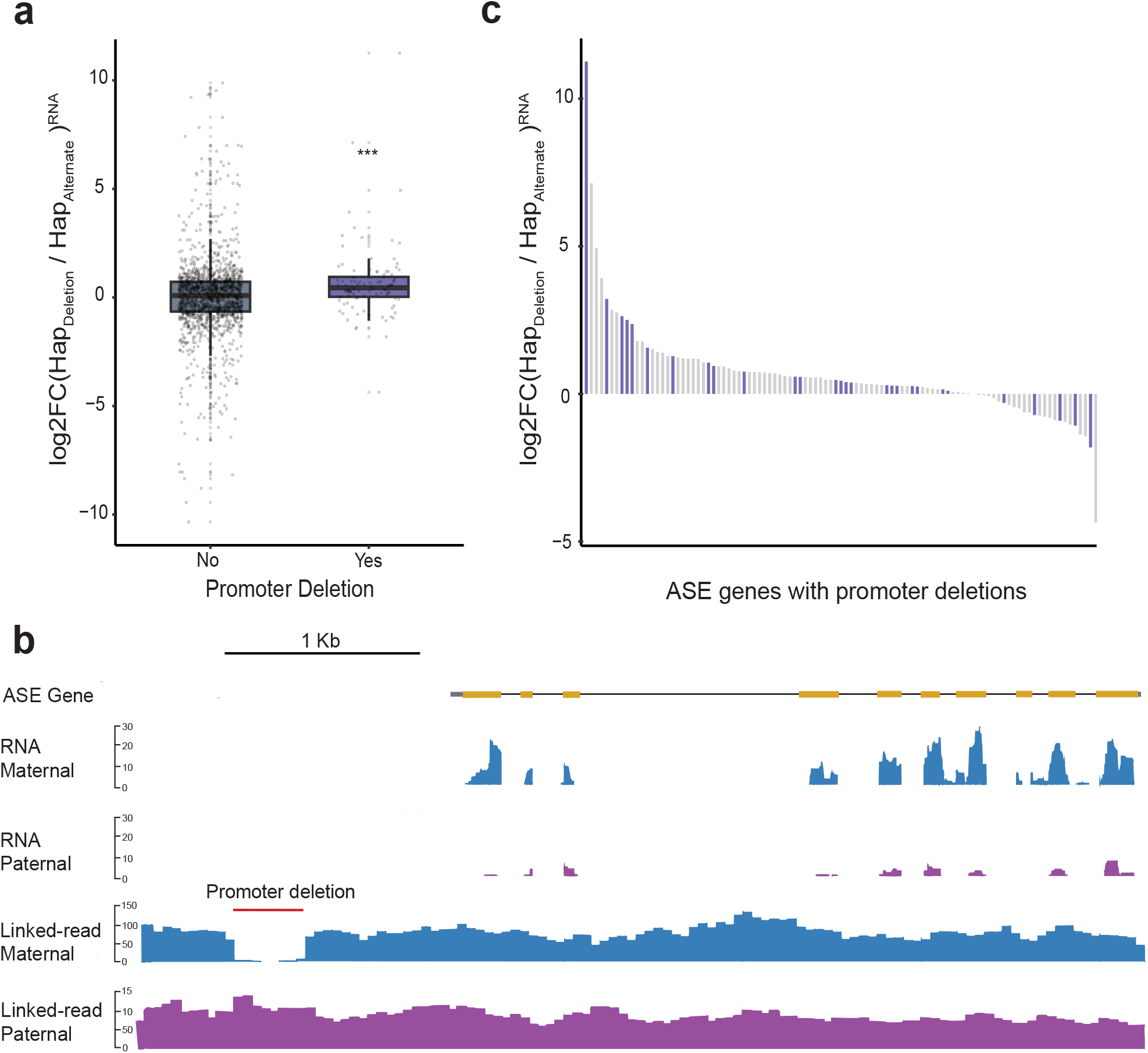
Structural variation as a potential source of cis-regulatory variation between haplotypes. **a)** 1,654 phased deletion:ASE gene pairs for which the gene and its nearest deletion are in the same phase block were identified. Comparison of allele-specific expression of genes with a deletion in their promoter (1 Kb upstream of TSS) (n = 104) versus those whose nearest deletion was not in its promoter (n = 1550). Gene expression is expressed as a ratio of read counts for the deletion-containing allele (HAP_Deletion_) relative to the alternate allele (HAP_Alternate_). For genes where there is no structural variant in the promoter, HAP_Deletion_ represents the allele in which the nearest deletion is found. Genes with a deletion in their promoter (n=104) had significantly higher expression of the deletion-containing allele (Hedges G effect-size 0.378) compared to ASE genes without promoter deletions (*** = 10,000 permutations, p < 0.001) **b)** Genome browser view of RNA-seq read coverage phased between maternal (purple) and paternal (blue) haplotypes for a predicted aldehyde dehydrogenase gene (FAIR_021608). Phased linked-reads between haplotypes reveal a 452 bp deletion (red bar) in the maternal allele that is associated with increased gene expression. **c)** A motif enrichment analysis of sequence absent in the ‘deleted’ allele of the 104 promoter:gene pairs from ASE genes identified four significantly enriched motifs, with the most abundant (n=28) matching the binding site of zinc-finger protein 1 (AZF1). ASE genes are ordered by their allelic expression ratio, and bars are colored based on whether the promoter deletion contains the AZF1 motif. There was no significant effect of AZF1 motif presence on allele-specific expression, but the sample size was small (n=28; 10,000 permutations, p = 0.2717).

### Allele-specific accessible chromatin regions (AS-ACRs) are linked to expression of proximal genes

CRMs can alter the magnitude, timing, and localization of transcription and divergence in CRM activity between haplotypes is a source of variation in ASE. We reasoned that differences in cis-regulatory module activity between haplotypes may be reflected in chromatin accessibility. Using the same framework for detecting ASE, we tested 5,423 ACRs with 2 or more polymorphisms for haplotype-specific differences in levels of chromatin accessibility. A total of 137 allele-specific ACRs were identified at a 5% false discovery rate (Benjamini Hochberg) and these were primarily classified as genic (58%), while 34% are proximal (within 2 Kb of nearest gene), and 11.6% are distal (greater than 2 Kb from nearest gene) (Additional File 1: Table S2). The epigenetic profile of AS-ACRs and their flanking sequence (2 Kb) was similar to all ACRs, including the subset of 5,423 ACRs with 2 or more polymorphisms used for haplotype-specific analysis (Additional File 2: Figure S12). Notably, only 2.5% of the 5,423 ACRs exhibited significant haplotype-specific chromatin accessibility, compared to 28% of genes with significant ASE. This may be a consequence of levels of sequencing coverage, where reduced coverage (i.e. for ATAC-seq) limits detection of differences in chromatin accessibility between alleles. It is also possible that genetic variants could disrupt transcription factor binding to impact allelic expression without affecting local chromatin structure.

Using locally phased blocks, we associated allele-specific differences in chromatin accessibility (AS-ACR) with allele-specific expression. Of the 137 AS-ACRs, 94.1% resided in the same phased block as their closest ASE gene with a median of 7,069 bp of intervening sequence. Next we correlated levels of chromatin accessibility of AS-ACRs with gene expression of ASE-genes on a per haplotype basis. For ACR-gene pairs separated by 5 Kb or less (n=51) there is a significant correlation between chromatin accessibility and gene expression (R = 0.69, p < 0.001) (Fig. 4a), with higher levels of allelic chromatin accessibility associated with an increase in expression of the proximal gene. The association between AS-ACRs and gene expression is reduced when examining ACR-gene pairs separated by more than 5 Kb (R = 0.46, p < 0.001) (Additional File 2: Figure S13). This pattern is consistent with the statistical modeling of ASE which highlights the importance of local chromatin state in predicting differences in gene expression between haplotypes (Fig. 2b).

**Fig 4:**
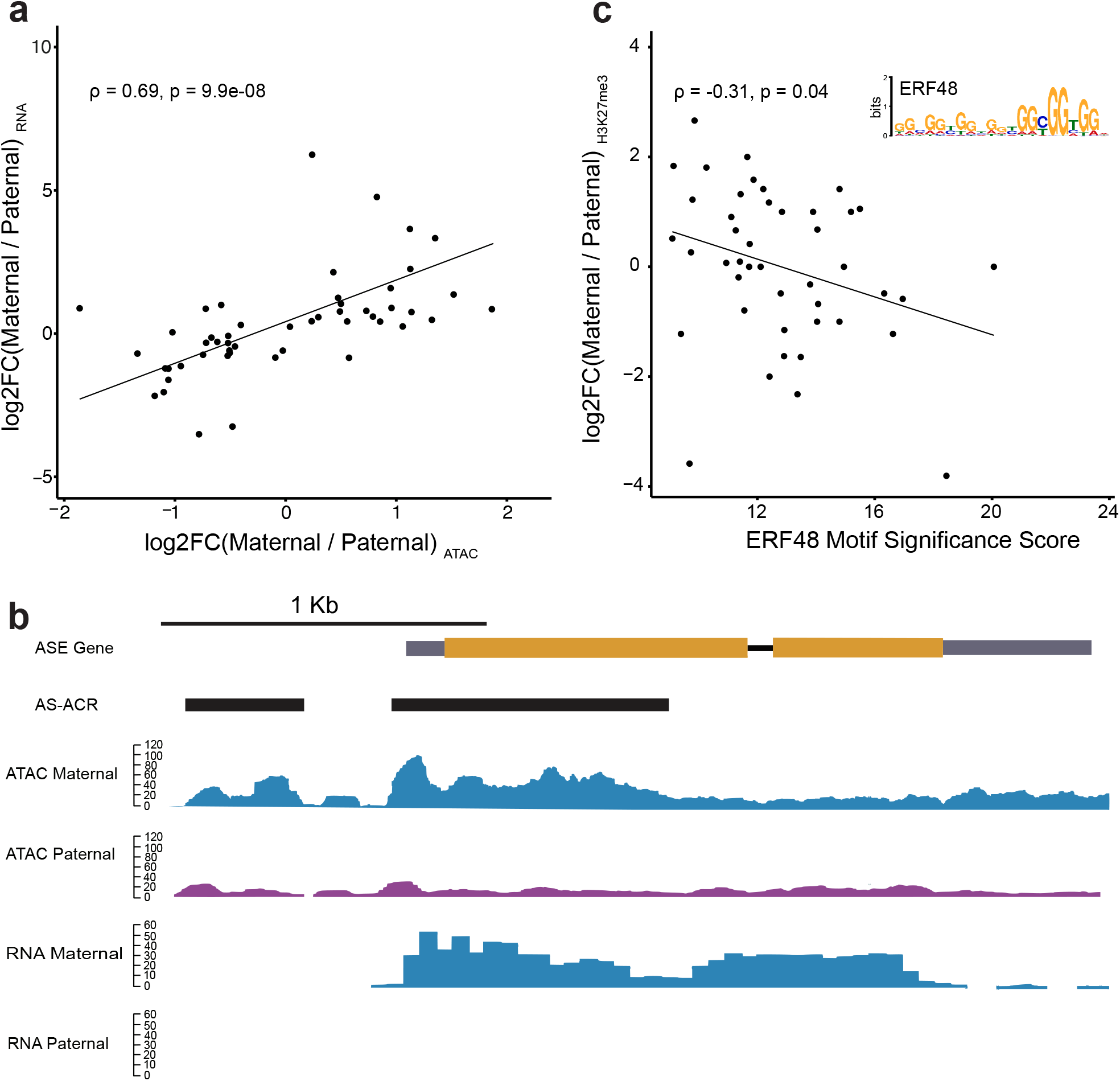
Allelic chromatin accessibility is linked to allele-specific expression of proximal genes. **a)** Allele-specific ACRs (AS-ACRs) were paired with their nearest ASE genes, restricting the analysis to ACR-gene pairs separated by 5 Kb or less (n=51) while residing within the same phase block. Allele-accessibility of AS-ACRs represented as the ratio of maternal: paternal ATAC-seq reads is positively correlated with allele-specific expression of neighboring genes within 5 Kb (n = 51, ρ = 0.69, p = 9.9e-08). **b)** Genome browser view of two AS-ACRs (black bars) located in the gene body and promoter of an ASE gene encoding a 7-deoxyloganetic acid glucosyltransferase (FAIR_019693). Monoallelic expression of the maternal allele (blue) is associated with increased chromatin accessibility of both AS-ACRs in the maternal haplotype. **c)** Motif enrichment analysis of the sequence underlying the paternal alleles of AS-ACRs identified the binding motif of ERF48 (see inset). Significant similarity of paternal sequences of AS-ACRs was observed for 44 / 137 AS-ACRs. The ERF48 motif significance score is correlated with allelic H3K27me3, where higher similarity of the paternal sequence to the canonical motif is associated with greater levels of H3K27me3 in the paternal haplotype.

A clear demonstration of the link between allelic chromatin accessibility and expression of proximal genes occurs at a gene predicted to encode a 7-deoxyloganetic acid glucosyltransferase (Additional File 1: Table S20). Two AS-ACRs were detected at this gene, one in the first exon, and the other in the gene promoter (Fig. 4b). The maternal alleles of the promoter AS-ACR and genic ACR had a log_2_FC (maternal / paternal ATAC) of 1.51 and 2.20, respectively. This coincides with monoallelic expression of the maternal allele, with no detectable expression of the paternal allele (Fig. 4b). The difference in accessibility in the gene body may be a consequence of active transcription of the maternal allele, while differences in accessibility of the promoter may reflect differential binding of transcription factors to the maternal and paternal alleles. Both the maternal and paternal sequence underlying the AS-ACR (364 bp) contains a hAT DNA transposon and a LTR transposon. Several studies have noted the role of transposable elements (TE) in generating variation in CRMs. This may occur through the disruption of functional cis-regulatory sequences by TE insertion [26,27], or TEs may contain cis-regulatory sequences with the potential to rewire gene regulatory networks by bringing novel CRMs in close proximity to genes [26,28]. DNA transposons, in particular, are a significant source of species-specific ACRs in angiosperms [19]. Similarly, AS-ACRs in the ‘Fairchild’ genome are enriched for DNA transposons relative to all detected ACRs (1,000 permutations, p = 0.002) (Additional File 2: Figure S14), with the hAT subfamily of class II DNA transposons being abundant in sequences underlying AS-ACRs. We did not observe any significant enrichment for class I transposons (Additional File 2: Fig. S15).

We searched the sequences underlying the maternal and paternal alleles of AS-ACRs for known transcription factor binding sites using MEME-ChIP [29]. Four motifs were enriched in either maternal or paternal alleles of AS-ACRs (Additional File 1: Table S21,S22). The motifs were highly repetitive in sequence which complicated the identification of specific transcription factor binding sites. One motif enriched in paternal alleles of AS-ACRs matches the ERF48 binding motif also known as DREB2C (Fig. 4c) (Additional File 1: Table S23). ERF48 binding motifs were also found to be enriched in differentially accessible chromatin regions between cotton (*Gossypium*) accessions before and after allopolyploidization, interspecific hybridization, and domestication [30], suggesting ERF48 motifs are a potential source of cis-regulatory variation across multiple plant species. ERF48 is a target of Polycomb repressive complex 2 which deposits H3K27me3 [30,31] and typically results in reduced transcriptional activity. Following this logic, we explored the relationship between levels of H3K27me3 at paternal alleles of AS-ACRs and the presence of the ERF48 motif. The ERF48 motif was detected in the paternal allele of 44 of the 137 AS-ACRs (Additional File 1: Table S23) and the degree of similarity to the consensus ERF48 motif for each AS-ACR correlated with allelic H3K27me3 (Spearman correlation, ρ = −0.31, p = 0.04) (Fig. 4c). Paternal haplotypes of AS-ACRs with greater sequence similarity to the ERF48 motif had greater levels of paternal H3K27me3 highlighting the potential for AS-ACRs to reveal haplotype-specific CRM that may be a consequence of differential activity of chromatin remodelers caused by sequence divergence between haplotypes.

## Discussion

Cis-regulatory variants are a major source of gene expression divergence between species [32,33], but unraveling the relationship between cis-regulatory variation and allele-specific differences in gene expression is challenging in the absence of haplotype-phased genome assemblies. In this study, we assembled a locally phased reference genome for ‘Fairchild’ mandarin and used it to investigate the molecular hallmarks of allele-specific expression. ‘Fairchild’ is a mandarin derived from parental cultivars with shared pummelo ancestry and, as a result, genes in these regions identical by state cannot be evaluated for allele-specific differences in gene expression or chromatin state. In cultivated mandarins with similar ancestry to ‘Fairchild’, these regions of homozygous pummelo ancestry account for 12-38% of the genome [4]. Still, genes in heterozygous portions of the genome could be used to explore the extent of allele-specific expression. We found that allele-specific expression is common, with differential expression between maternal and paternal alleles occuring at nearly one third of the 7,300 genes with two or more genetic variants. We expect that even more genes with ASE would be detected in hybrids with fewer regions of shared ancestry.

Focusing on the set of 2,071 ASE genes, we investigated chromatin features associated with differential transcription of alleles. To do so, we integrated data sets for gene expression, chromatin accessibility, and histone modifications (among others). We found that 30.69% of variation in allele-specific expression can be inferred from local chromatin state, with the most influential factors located in the gene region, as opposed to promoter or other distal regulatory sequences (Fig. 2b). It is important to note that our analysis has revealed associations between allelic chromatin dynamics and gene expression. Without genome-wide surveys of chromatin in relevant mutants or directed manipulation of the chromatin state in regions of interest, it is not possible to determine whether allele-specific histone modifications are driving differential gene expression or if they are a consequence of differential transcription. In ‘Fairchild’, the factors most associated with allele-specific expression were chromatin accessibility and histone modifications in gene regions, including H3K36me3, H3K56ac, and H3K27me3. The next most influential set of factors were those located in promoter regions, including haplotype-specific accessibility of chromatin and the presence or absence of a structural variant. In contrast to genic and promoter regions, the effect of factors in distal regulatory regions was relatively limited, although two factors (ATAC, H3K27me3) were still identified as having significant effects. Distal regulatory regions with functional roles have been identified in large-genome species including maize and wheat [15,34], but distal ACRs seem to be less common in smaller genomes [15,35,36]. In citrus, we expect fewer distal ACRs since the average intergene distance in the ‘Fairchild’ genome is around 2.7 Kb. In fact, only 17.3% of ACRs detected in our dataset were classified as distal (Additional File 1; Table S2). Additionally, the correlation of AS-ACR chromatin accessibility and ASE decays for ACR-gene pairs that are separated by more than 5 Kb (Additional File 2: Figure S17). Here, we restricted analysis of chromatin accessibility and gene expression to neighboring ACR-gene pairs with no intervening protein-coding gene sequence. In other angiosperms, there are typically no intervening genes between ACRs and their target gene even in large-genome species like maize, where pairs are separated by many tens of kilobases [15,19]. While this is a reasonable assumption, future integration of conserved non-coding sequences, chromatin contact maps, and tissue-specific identification of ACRs will enable a more complete characterization of the distal regulatory landscape in small genome species like citrus to more precisely quantify the contribution of these sequences to gene expression regulation.

Next, we sought to understand and characterize potential sources of cis-regulatory variation in ‘Fairchild’. Structural variation between haplotypes is one possible source of cis-regulatory variation and may alter TF binding affinity, introduce new cis-regulatory modules, or perturb the coordination of neighboring cis-regulatory elements. Genome-wide surveys of genetic variation in crop species have revealed extensive levels of structural variation [6,9,37,38]. In some cases, these variants have been linked to functional differences [5,39,40]. In cultivated tomato and its wild relatives, for example, 7.3% of genes with structural variants in promoter regions were differentially expressed [8]. In ‘Fairchild’, structural variants are enriched in the promoters of ASE genes, with associated sequences predominantly derived from hAT and MULE-MuDR DNA transposons. The designation of ‘insertion’ or ‘deletion’ is relative to the ‘Fairchild’ consensus sequence. Structural variants were not polarized to an outgroup species to determine their ancestral state. Haplotypes with the non-consensus allele are termed ‘deletions’ here and deletions in the promoters of ASE genes were associated with increased expression (Fig. 3a). The sequence underlying the longer, consensus ‘inserted’ allele was enriched for transposon associated sequences, suggesting that the ‘inserted’ allele is actually linked to reduced expression, possibly due to the associated TE. TE insertions may alter gene expression by disrupting cis-regulatory modules [41], or through the spreading of repressive chromatin modifications to nearby regulatory sequences [42,43]. While TE-mediated disruption of local sequence is likely a source of SV-related ASE, our analysis also reveals evidence for a signature of TE-associated CRM with repressive effects on gene expression. The prevalence of SV-related cis-regulatory variants in ‘Fairchild’ suggests that the comparison of chromatin accessibility across structurally diverse citrus genomes is likely to further our understanding of the factors underlying gene expression divergence in heterozygous perennial tree crops species, like those in the genus *Citrus*.

### Genetic variation in CRMs is another potential source of allele-specific expression

Genetic variants that perturb transcription factor binding may subsequently alter local chromatin remodeling. One example of this is the pioneer TF LEAFY which binds to nucleosome occupied binding sites at the *AP1* locus and recruits SWI/SNF chromatin remodelers [44]. This is followed by increased chromatin accessibility and elevated levels of *AP1* expression [44]. We explored the sequences underlying AS-ACRs to identify genetic variants with possible impacts on TF binding and subsequent gene expression. The sequences underlying AS-ACRs were enriched for DNA transposons. In genome-wide surveys of chromatin accessibility across many angiosperm species, DNA transposons were also enriched in a set of species-specific ACRs [19]. To begin to understand how genetic variation at putative CRMs impacts phenotype, we mined ACRs with haplotype-specific patterns of accessibility for candidate transcription factor binding motifs which may underlie the differential gene expression patterns observed in this citrus hybrid. We detected the enrichment of ERF48 binding sites in paternal alleles of AS-ACRs, and observed a correlation between ERF48 motif presence and allelic H3K27me3 (Fig. 4c). ERF48 binding sites were also found to be enriched in differential DNAse hypersensitive regions discovered in a panel of cotton accessions spanning polyploidization and domestication in the species [30]. Han and colleagues speculate that PRC2 binds the guanine-rich binding site of ERF48, and that PRC2-mediated deposition of H3K27me3 may be involved in transitions between states of chromatin accessibility [30]. The sequence-specific DNA binding activity of H3K27me3 “writers” such as Polycomb repressive complex 2 (PRC2), and “erasers” such as JUMONJI DOMAIN-CONTAINING PROTEINS (JMJ) may underlie differential regulation of alleles with resulting ASE. An extensive yeast-one hybrid screen revealed 233 transcription factors that bound motifs associated with PRC2 binding and subsequent H3K27me3 deposition [45]. Similarly, the sequence-specific DNA binding activity of JMJ proteins such as RELATIVE OF EARLY FLOWERING 6 (REF6) [46–48] suggests that sequence alterations in its binding motif could produce differential H3K27me3 demethylation between alleles. We find a significant correlation between ERF48 motif similarity scores and allelic levels of H3K27me3, but no significant relationship between allelic chromatin accessibility and allelic levels of H3K27me3 of AS-ACRs, but we are likely limited by the small sample size of AS-ACRs, the sequence depth of ATAC-seq libraries, and the conservative method used for ACR identification. Importantly, AS-ACRs are indistinguishable from other ACRs when examining ATAC-seq coverage without read-phasing (Additional File 2: Fig. S12). ERF48 motif enrichment in AS-ACRs highlights an instance of how sequence variation in a TF binding site relates to local chromatin dynamics and gene expression variation.

Further mining ACRs for functional variants is likely to reveal genotype-phenotype associations. In maize, 40% of genetic variants with heritable effects of phenotypes were located in the 1% of the genome marked by accessible chromatin [49]. In ‘Fairchild’, approximately 2% of the genome resides in ACRs and, similar to maize, we expect these sequences to be enriched for functional variants. The functional consequences of cis-regulatory variation in citrus have previously been demonstrated for two important commercial traits [5,6]. Allele-specific expression is an indication that there are functional cis-acting sequence variants. The 2,071 genes with ASE in ‘Fairchild’ are good candidates for exploring the phenotypic consequences of cis-regulatory sequence variation. We highlight two of these genes as candidates for further study. One AS-ACR was discovered upstream of a gene encoding a 7-deoxyloganetic acid glucosyltransferase (Fig. 4b) which catalyzes the production of an iridoid monoterpene [50]. Iridoid monoterpenes have been shown to be involved in plant-insect interactions and are of interest as natural products because of their pharmacological activities [51–53]. A second gene of interest is one predicted to encode a homolog of potato SNAKIN-1 (Additional File 1: Table S24). A 165 bp structural variant was identified in the promoter of this gene in ‘Fairchild’. SNAKIN-1 is a small peptide identified in potato, that has demonstrated antimicrobial activity against both bacterial and fungal pathogens when heterologously expressed in several plant species [54–57]. The role of small antimicrobial peptides has recently been highlighted for their potential in combating citrus greening disease [58]. These are only two examples where connections between genetic variation, chromatin accessibility, and allele-specific differences in gene expression may lead to functional phenotypic differences. Further dissection of these loci and others with evidence of allele-specific expression will likely reveal troves of variation relevant for genetic improvement of cultivated citrus.

## Conclusions

We assembled a locally phased reference genome for ‘Fairchild’ mandarin and used it to investigate the molecular signatures of allele-specific expression. By profiling gene-expression, chromatin accessibility, DNA methylation, and several histone-modifications we provide a detailed view of gene regulation in this hybrid cultivar. We document the extent of haplotype-specific differences in gene regulation and begin to dissect the interplay between local variation in chromatin dynamics and gene expression. We find examples where allele-specific expression is associated with local structural variation in promoter sequences and others where expression differences are more likely due to genetic variation in putative TF binding. We anticipate that the datasets produced here will inform approaches for studying the interplay between genetic variants in heterozygous genomes and molecular phenotypes such as gene expression, chromatin accessibility, and epigenetic modifications with the goal of revealing functional cis-regulatory sequences and understanding the evolution of gene regulation.

## Materials and methods

### Sampling

Fairchild mandarin was sampled from the Givaudan Citrus Variety Collection at the University of California Riverside. Young leaves were collected from a greenhouse grown individual of the cultivar ‘Fairchild’ mandarin (*Citrus reticulata*) (Accession No: PI 539508, Inventory No: RIV 2014 PL). To deplete starch in the leaves, the tree was covered with brown paper bags 24 hours prior to sampling. The leaves were sampled early in the morning and immediately frozen in liquid nitrogen. The intact, frozen leaves were then shipped to Arizona Genomics Institute (AGI) for DNA extraction from nuclei for PacBio sequencing and to the Amplicon Express company (Pullman, WA) for DNA extraction for Illumina, Bionano, and 10X sequencing.

### DNA extraction and sequencing

High-molecular-weight (HMW) DNA was extracted and used for the library construction. SMRTBell libraries (20 Kb) were prepared according to the manufacturer’s protocol including gTube size fraction and BluePippin size exclusion of fragments smaller than 20kb before constructing libraries for SMRTbell template construction. Sequencing on the Pacbio RS II machine was performed at AGI following library construction. A total of 3,930,477 reads were produced from 53 flow cells (69 Gb of sequence). Sequenced reads had an N50 of 15 Kb and a maximum length of 78 Kb. For scaffolding, a 10X sequencing library was constructed according to the 10X Genomics protocol (Pleasanton CA USA) by Amplicon Express. This library was then sequenced on the Illumina HiSeq4000 system producing 285 million 2 x 150 bp reads.

### Bionano genome (BNG) optical map construction and data analysis

High-molecular weight (HMW) DNA was isolated by the company Amplicon Express (Pullman, WA). The nicking enzyme Nb.BssS1 was chosen based on the de novo draft assembly. The high-quality HMW DNA molecules were treated with Nb.BssS1 (New England BioLabs, Ipswich, MA) to generate single-strand breaks with sequence-specific motifs (GCTCTTC). The nicked DNA molecules were then stained using the IrysPrep Reagent Kit (Bionano Genomics, San Diego, CA) and loaded onto the nanochannel array of IrysChip (Bionano Genomics) following the manufacturer’s guidelines. Imaging and data analysis were performed on the Irys system (Bionano Genomics). DNA molecules larger than 100 Kb were processed into BNX files using the software AutoDetect (v2.1.4) and only molecules of at least 180 Kb were assembled into the BNG map with the Bionano Genomics assembly pipeline [59,60], The p-value thresholds used for pairwise assembly, extension/refinement, and final refinement stages were 2×10 ^-8^, 1×10 ^-9^, and 1×10 ^-15^, respectively. The de novo assembly was digested in silico at Nb.BssS1 restriction sites using Knickers and aligned with the BNG map using RefAligner in Bionano Solve (v03062017). Visualization of the alignments were performed with IrysView.

### Genome assembly

We *de novo* assembled the ‘Fairchild’ genome using PacBio long reads (RSII) with a diploid assembler, FALCON (v1.8.6). In total, 69 Gb of PacBio long reads were generated from 53 flow cells (Additional File 3: Table S1), which is roughly equivalent to 187 fold of the estimated genome size (365 Mb). An initial assembly generated a 429 Mb primary genome sequence with contig N50 of 2.3 Mb and 69 Mb of secondary sequences with a contig N50 of 144 Kb. The assembly errors were corrected with Quiver (v2.0) by making a consensus sequence from alignments of the raw PacBio long reads mapped to the de novo assembly. A second round of polishing was performed with Pilon (v1.20) based on the alignment of 150 bp paired-end Illumina short reads from sequencing of 10X Linked-Reads. To build a haploid reference genome, we used Haplomerge2 (v11242015) to remove redundancy and merge overlapping contigs in the assembly, which resulted in a haploid genome of 360 Mb with contig N50 of 3.6 Mb. The BioNano optical map data were assembled with RefAligner in Bionano Solve (v03062017). We aligned the BioNano optical map contigs with PacBio contigs using HybridScaffold.pl in Bionano Solve (v03062017) to correct mis-assemblies and further scaffold contigs. We then assembled 10x linked reads based on the barcode of each single molecule and filled gaps in scaffolds with a graph-based method (Du et al, 2017). These high-quality scaffolds were then anchored to chromosomes by integrating three genetic maps (reference) and synteny between citrus genomes as implemented in ALLMAPS (v07222015). The final scaffolded genome was 365 Mb in length and 94.8% of contigs (346/365 Mb) were anchored to nine chromosomes. (Additional File 3: Tables S2, S3) We assessed the completeness of the genome assembly using BUSCO (v5.5.0) against the embryophyta_odb10 dataset and the BUSCO completeness was estimated to be 98.5 %. LTR index (LAI) was calculated with EDTA (v1.7.0) and reported a LAI of 15, indicating a highly continuous assembly of intergenic regions.

### RNA-seq for allele-specific expression

RNA-seq was performed as previously described [19]. Leaves were flash-frozen with liquid nitrogen immediately after collection. The samples were ground using a mortar and pestle. Total RNA was extracted and purified using TRIzol Reagent (Thermo Fisher Scientific) following the manufacturer’s instructions. For each sample, 1.3 μg of total RNA was prepared for sequencing using the Illumina TruSeq mRNA Stranded Library Kit (Illumina) following the manufacturer’s instructions.

### ATAC-seq

ATAC-seq was performed as described previously [61]. Approximately 200 mg of flash-frozen leaves were immediately chopped with a razor blade in approximately 1 ml of pre-chilled lysis buffer (15 mM Tris-HCl pH 7.5, 20 mM NaCl, 80 mM KCl, 0.5 mM spermine, 5 mM 2-mercaptoethanol and 0.2% Triton X-100). The chopped slurry was filtered twice through Miracloth and once through a 40 μm filter. The crude nuclei were stained with 4,6-diamidino-2-phenylindole (DAPI) and loaded into a flow cytometer (Beckman Coulter, MoFlo XDP). Nuclei were purified by flow sorting and washed as described previously [61]. The sorted nuclei were incubated with 2 μl Tn5 transposomes in a 40 μl tagmentation buffer (10 mM TAPS-NaOH pH 8.0, 5 mM MgCl2) at 37 °C for 30 min without rotation. The integration products were purified using a Qiagen MinElute PCR Purification Kit or NEB Monarch DNA Cleanup Kit and then amplified using Phusion DNA polymerase for 10–13 cycles. PCR cycles were determined as described previously [18]. Amplified libraries were purified using AMPure beads to remove primers.

### ChIP-seq

Chromatin immunoprecipitation (ChIP) was performed using a previously described protocol [62]. In brief, approximately 1 g freshly harvested leaves or flash-frozen leaves was chopped into 0.5 mm cross-sections and cross linked as described previously [62]. The samples were immediately flash-frozen in liquid nitrogen after crosslinking. Nuclei were extracted and lysed in 300 μl of lysis buffer. Lysed-nuclei suspension was sonicated using a Diagenode Bioruptor on the high setting, 30 cycles of 30 s on and 30 s off. To make antibody-coated beads, 25 μl Dynabeads Protein A (Thermo Fisher Scientific, 10002D) was washed with ChIP dilution buffer and then incubated with around 1.5 μg antibodies (anti-H3K4me3, Millipore-Sigma, 07–473; anti-H3K36me3, Abcam, ab9050; anti-H3K27me3, Millipore-Sigma, 07–449; anti-H3K56ac, Millipore-Sigma, 07–667) in 100 μl ChIP dilution buffer for at least 1 h at 4 °C. The sonicated chromatin samples were centrifuged at 12,000 g for 5 min and the supernatants were diluted tenfold in ChIP dilution buffer to reduce the SDS concentration to 0.1%. ChIP input aliquots were collected from the supernatants. For all of the samples and replicates, 300–500 μl of diluted chromatin was incubated with the antibody-coated beads at 4 °C for at least 4 h or overnight, then washed, reverse-crosslinked and treated with proteinase K in accordance with the protocol [62]. DNA was purified using a standard phenol–chloroform extraction method, followed by ethanol precipitation. The DNA samples were end-repaired using the End-It DNA End-Repair Kit (Epicentre) following the manufacturer’s protocol. DNA was cleaned up on AMPure beads (Beckman Coulter) with size selection of 100 bp and larger. The samples were eluted into 43 μl Tris-HCl and subsequently underwent a 50 μl A-tailing reaction in NEBNext dA-tailing buffer with Klenow fragment (3′ −> 5′ exo−) at 37 °C for 30 min. A-tailed fragments were ligated to Illumina TruSeq adapters and purified with AMPure beads. The fragments were amplified using Phusion polymerase in a 50 μl reaction following the manufacturer’s instructions. The following PCR program was used: 95 °C for 2 min; 98 °C for 30 s; then 15 cycles of 98 °C for 15 s, 60 °C for 30 s and 72 °C for 4 min; and once at 72 °C for 10 min. The PCR products were purified using AMPure beads to remove primers.

### MethylC-seq

Several leaves were immediately flash-frozen after harvesting and ground to a powder in liquid nitrogen. DNA was extracted and purified with the DNeasy Plant Mini Kit (Qiagen) and 130 ng were used for MethylC-seq library preparation. MethylC-seq libraries were prepared using a previously described protocol [63]; however, we used a final PCR amplification of eight cycles.

### Annotation of repetitive elements

To annotate and remove repetitive regions from the gene prediction process,, we generated a library of repetitive elements with RepeatModeler v. 2.0.1 [64] using default parameters. This library was used to soft-mask these elements in the genome with RepeatMasker v. 4.1.1 [65] (Additional File 3: Table S4). We used the calcDivergenceFromAlign.pl script, available in the util directory of RepeatMasker package, to calculate and summarize the divergence from the consensus for each predicted repeat using the Kimura 2-Parameter distance with an adjustment for higher mutation rates at CpG sites. The createRepeatLandscape.pl script was then used to create a repeat landscape plot and visualize the repeat elements diversity, their relative age and divergence, using the divergence summary data (Additional File 2: Figure S18).

### Genome annotation

RNA-seq data was generated from young leaves and young flowers of ‘Fairchild’. A total of 33.4 Gb of sequence was produced on the Illumina HiSeq4000 (2 x 150 bp reads). Samples were collected and then flash frozen in liquid nitrogen and stored at −80°C. Total RNAs were extracted from these samples using TRIzol reagent (Thermo Fisher Scientific). RNA-Seq libraries were prepared using NEBNext Ultra Directional RNA Library Prep Kit. Libraries were prepared using an NEB rRNA depletion kit. Genome annotation was conducted with the Funannotate pipeline v. 1.8.10 [66]. First, the RNAsequencing (RNA-seq) data were assembled using Trinity v. 2.11.0 and aligned with PASA v. 2.4.1 to train the ab initio gene predictors [67–69]. After training the predictors Augustus v. 3.3.3 and SNAP v. 2013_11_29 with the evidence, a collection of gene predictors were run CodingQuarry v. 2.0, Augustus v. 3.3.3, GeneMark-ETS v. 4.62, GlimmerHMM v. 3.0.4, and SNAP v. 2013_11_29 [70–74]. These predictions were combined into a consensus gene model set with EVidenceModeler v. 1.1.1 [69]. Transfer rRNA genes were predicted using tRNAscan v. 2.0.9 [75]. Next, using HMMER v. 3 [76] or diamond BLASTP v. 2.0.8 [77,78] protein annotations and gene products were predicted for the consensus gene models based on similarity to Pfam [79], CAZyme domains with dbCAN HMM profiles [80–83], MEROPS [84], eggNOG v. 5.0 [85], InterProScan v. 5.51.85.0 [86], and Swiss-Prot [87] databases. In addition, Phobius [88] and SignalP v.5.0b [89] were used to identify transmembrane proteins and secreted proteins. (Additional File 3: Table S5)

### Variant calling and haplotype phasing

Single nucleotide polymorphisms (SNPs), small indels (< 41 bp), and structural variants were identified on the nine primary chromosomes using the 10x Genomics Longranger pipeline (10x Genomics, Pleasanton California). In short, 10x Genomics linked reads were aligned to the reference genome using the aligner Lariat which is based on the Random Field Aligner algorithm [90] https://github.com/10XGenomics/lariat. Variants were subsequently identified using GATK Haplotypecaller and GenotypeGVCF following the current GATK best practices [91]. In Longranger, insertions and deletions ranging from 40 bp - 30 Kb were detected using haplotype-specific reductions in read coverage, discordant read pairs, and local re-assembly. SNPs, indels, and SVs were then phased by aligning the linked-reads to the sequence of each allele to define which haplotype the read supports. A total of 1,015 phase blocks were constructed using 2.3 million phased heterozygous variants (mean length of phase block = 317,484 bp; standard deviation = 711,178.7 bp). Within each phase block, SNPs are attributed to either haplotype 1 or haplotype 2. When comparing SNPs between phase blocks, the assignment of haplotype 1 or 2 is random and not associated with an assignment of ancestry. Biallelic SNPs were identified and filtered using the following criteria with bcftools [92]: QD > 2.0 && FS < 60.0 && MQ > 40.0 && MQRankSum > −12.5 && N_ALT = 1 && N_MISSING = 0 && QUAL > 500 && MIN(FMT/DP) > 20 && INFO/DP < 450. This yielded 1.53 million phased heterozygous variants that were used in downstream analyses. The majority of variants are SNPs (1.3 million) that were used to phase sequencing reads for whole-genome re-sequencing, RNA-seq, ATAC-seq, bisulfite-seq, and ChIP-seq (H3K4me3, H3K36me3, H3K56ac, H3K27me3).

### Assignment of ancestry to phased SNPs

Whole-genome re-sequencing reads from maternal parent clementine mandarin [4], paternal parent ‘Orland’, and hybrid ‘Fairchild’ were trimmed for quality and sequencing adapter removal using Trimmomatic 0.36 (SLIDINGWINDOW:4:20 MINLEN:50 ILLUMINACLIP:LEADING:3 TRAILING:3) [93]. Reads were aligned to the ‘Fairchild’ reference genome using BWA mem 0.7.17 [94]. PCR duplicates were filtered using Picard MarkDuplicates [95]. Variants were identified using GATK Haplotypecaller and GenotypeGVCF following the current GATK best practices [91]. High-quality biallelic SNPs were identified and filtered using the following criteria with bcftools [92]: QD > 2.0 && FS < 60.0 && MQ > 55.0 && SOR < 3 && MQRankSum > −12.5 && ReadPosRankSum > −2.0 && QUAL > 500 && MIN(FMT/DP) > 30 && INFO/DP < 450. The remaining 1.3 million variants were used in combination with whole-genome sequencing reads from both parents to perform pedigree-based phasing using Whatshap v. 1.17 [96]. A constant recombination rate of 3.0 cM/Mb (average observed in *C. clementina* [97] was used for pedigree-based phasing of SNPs. After ancestry assignment, SNPs heterozygous in one or both parents were removed, leaving only SNPs that are homozygous for alternate alleles in the two parents (141,290 SNPs). The ancestry assignment of these SNPs was combined with their read-based haplotype assignment in order to impute the ancestry of all 1.3 million SNPs between the two haplotypes. In total, 394 phase blocks contained high quality SNPs and were designated as unclassified, non-recombinant, or recombinant depending on the ancestry composition of SNPs within the phase block (Additional File 1: Table S10; Additional File 3: Table S6). Unclassified phase blocks (n = 149) did not contain any SNPs that could be assigned ancestry, for non-recombinant phase blocks (n= 231) the ancestry of every SNP was consistent within each haplotype of the phase block, in recombinant blocks (n=14) the ancestry of SNPs was variable within each haplotype of the block. For non-recombinant blocks, the ancestry of SNPs that could not be assigned using parental genotypes was imputed using the ancestry of SNPs within the same phase block. For recombinant blocks, only SNPs that could be assigned ancestry using parental genotypes were retained. One recombinant phase block with repeated haplotype switching of SNP ancestry was removed from the analysis (2983 SNPs). Additionally, SNPs whose haplotype ancestry was discordant with the majority of other SNPs in the phase blocks were filtered. The ancestry of SVs was imputed following this same procedure, with the exception of SVs in recombinant phase blocks. For SVs in recombinant phase blocks, the ancestry of the SV was imputed using the majority ancestry of the nearest 5 SNPs upstream and downstream of the SV.

### Polarization of heterozygous SNPs in ‘Fairchild’

Whole genome sequencing reads from *Poncirus trifoliata* accession DPI 50-7 (NCBI PRJNA648176) [98] were trimmed for quality and sequencing adapter removal using Trimmomatic 0.36 (SLIDINGWINDOW:4:20 MINLEN:50 ILLUMINACLIP:LEADING:3 TRAILING:3) [93]. Reads were aligned to the ‘Fairchild’ reference genome using BWA mem 0.7.17 [94]. PCR duplicates were filtered using Picard MarkDuplicates [95]. Variants were identified using GATK Haplotypecaller and GenotypeGVCF following the current GATK best practices [91]. High-quality biallelic SNPs that are homozygous in DPI 50-7 were identified and filtered using the following criteria with bcftools [92]: QD > 2.0 && FS < 60.0 && MQ > 55.0 && SOR < 3 && MQRankSum > −12.5 && ReadPosRankSum > −2.0 && QUAL > 500 && MIN(FMT/DP) > 30 && INFO/DP < 450. Next, these variants were intersected with heterozygous SNPs in ‘Fairchild’ and biallelic SNPs were retained. This final set of SNPs represent ‘Fairchild’ SNPs for which the ancestral state is indicated by the allele present in *Poncirus trifoliata* (Additional File 3: Table S7).

### Processing of ChIP-seq and ATAC-seq sequencing reads

Prior to alignment, ChIP-seq reads for four histone modifications (H3K4me3, H3K36me3, H3K56ac, H3K27me3) and ATAC-seq reads were trimmed for quality (SLIDINGWINDOW:4:15 MINLEN:50 ILLUMINACLIP:LEADING:3 TRAILING:3) and adapter sequences removed using Trimmomatic 0.36 [93]. Sequence reads were then aligned to the ‘Fairchild’ reference genome using BWA mem 0.7.17 [94]. PCR duplicates were filtered using Picard MarkDuplicates [95]. A summary of sequence reads for each data set before and after quality control is provided (Additional File 3: Table S8). Peak identification for histone modifications (ChIP-seq) and accessible chromatin (ATAC-seq) was performed using Genrich (https://github.com/jsh58/Genrich) and MACS2 [99] following [19]. Significant peaks within 100 bp were merged. Sequences underlying ACRs were aligned to plastid (NC_034671) and mitochondrial (NC_037463.1) genomes of *C. reticulata* using NCBI BLAST and significant hits were removed. The number of peaks identified, along with their average read depth and percentage of the genome occupied by these peaks is summarized here (Additional File 1: Table S1). Two methods were used for identifying accessible chromatin regions (ACRs). One method (MACS2) identified a total of 31,191 ACRs spanning 10.4 Mb with a fraction of reads in peaks (FRiP) score of 0.16, and the other (Genrich) identified 9,172 ACRs spanning 7.3 Mb (FRiP = 0.13) (Additional File 1: Tables S1, S2). There was 70% overlap between ACRs identified by Genrich and ACRs identified by MACS2. Additionally, 97% of Genrich ACRs overlapped ACRs identified with MACS2. An unpublished benchmarking of the ATAC-seq peak calling programs HMMRATAC, MACS2, and Genrich found that Genrich had the highest precision, recall, and F1 score [100]. For this reason we decided to use the ACRs detected with Genrich.

### Processing of bisulfite sequencing reads and analysis of DNA methylation

Bisulfite-sequencing reads were trimmed for quality (SLIDINGWINDOW:4:15 MINLEN:50 ILLUMINACLIP:LEADING:3 TRAILING:3) and adapter sequences were removed using Trimmomatic 0.36 [93]. Reads were then aligned to the ‘Fairchild’ reference genome using BatMeth2 [101] and PCR duplicates were removed using Picard MarkDuplicates [95]. Bisulfite-sequencing reads were phased between parental haplotypes with ancestry-assigned SNPs using Whatshap [96]. DNA methylation of cytosines was counted for both haplotypes separately using DNMtools [102] (Additional File 3: Tables S9, S10 and S11). The probability of differential methylation between haplotypes was determined using a one-directional version of Fisher’s exact test implemented in DNMtools [102,103]. Differentially methylated cytosines with read coverage >= 10 and probability of differential methylation >=0.7 were retained for modeling of ASE. Overall methylation was calculated for cytosines with coverage >= 10 using the full, unphased bisulfite-sequencing data with DNMtools [102]. Methylation status is based on the number of reads that support a C-to-T bisulfite conversion versus the total number of reads aligning to a given cytosine.

### Regional analysis of genomic features

Metaplots were generated to summarize regional patterns of chromatin levels, typically centered on the peak of an open chromatin region, as determined by ATAC-seq. For each data set, read coverage was scaled from 0 to 1 by dividing the read coverage at each site by the 98th quantile of sequencing coverage per sample [19]. ACRs and genes were scaled to a 1000 bp region, as well as their 2kb flanking regions using deepTools v.3.5.1 [104] and signal densities for each feature were plotted across these regions in 50 bp windows.

### Determination of allele-specific gene expression and chromatin accessibility

Read alignments for quality and adapter trimmed ATAC-seq and ChIP-seq were performed using BWA-mem 0.7.17 [94] as described above. RNA-seq reads were also quality and adapter trimmed using Trimmomatic 0.36 [93] prior to read mapping to the ‘Fairchild’ reference genome with the slice-aware alignment tool STAR 2.7.10 [105]. After initial alignment of RNA-seq, ATAC-seq, and ChIP-seq reads, WASP [106] was used to correct read-mapping bias (Additional File 3: Table S8). This is done by switching the reference allele to the alternate allele (and vice versa) in reads that align across SNPs. Sequence reads that do not align to the same genomic location are filtered to remove reads that cannot be definitively assigned to a haplotype. For each data set, allele-specific read counts were determined using GATK ASEReadCounter [91]. Allele counts were then intersected with genes (for RNA-seq) or peak regions (ATAC-seq and ChIP-seq). SNPs covered by less than 10 reads were removed. Genes and ATAC-seq peak regions with 2 or more SNPs were considered for subsequent allele-specific analysis. GeneiASE [107] was used to detect significant differences in the level of gene expression or chromatin accessibility between alleles. GeneiASE first measures individual SNP effects based on the read counts of each allele at a given SNP. The median SNP effect-size for genes with significant ASE was 4.6, while genes without significant ASE had a median SNP effect-size of 1.9. Next, a gene/peak test statistic is calculated by pooling all of the SNP effects that coincide with the given gene or peak using Stoufer’s method [108]. The significance of gene test-statistics was then determined by comparison to a null distribution of gene test statistics. This null distribution is characterized by an overdispersion parameter and mean marginal probability of success [107]. These parameters were determined by fitting a beta binomial model to whole-genome sequencing read counts overlapping the same gene/peak regions of interest. This provides an additional correction for read-mapping biases. Genes or peaks with significant allele-specific differences were identified at a false discovery rate of 0.05 (Benjamini-Hochberg) (Additional File 1: Table S7,S8; Additional File 3: Table S12).

### Integration of allele-specific ACRs (AS-ACRs) and ASE using phased blocks

Genes with significant ASE in both RNA-seq replicates and ACRs with significant allele-specificity were paired using phase block information. If an ASE gene and its nearest AS-ACR were located in the same phase block, the sum of read counts for the ASE gene (RNA-seq), and the AS-ACR (ATAC) was calculated for each haplotype. In this case, the read count of the maternal haplotype for the ATAC peak region corresponds to the level of expression of the maternal allele of the linked gene. The log_2_ fold change between the read counts of maternal and paternal haplotypes were determined both for gene expression and chromatin accessibility. A pseudo count of 1 was added to each value prior to log transformation.

### Permutation tests for genomic feature enrichment

A permutation test was performed to examine the enrichment of allele-specific ACRs for transposable element sequences. The number of AS-ACR-TE intersections was compared to a set of similarly sized regions selected from the complete set of ACRs and the frequency that each sample intersected annotated TEs. This process was repeated (n=1000). This analysis was performed for all TEs, and then for TEs separated into class I and class II transposons. A similar approach was used to examine the enrichment of structural variants (50 bp - 30 Kb in size) within promoters of ASE genes. In this case, the number of SV-promoter intersections was compared to a sample of similarly sized regions from promoters of genes analyzed for ASE. The R package regioneR was used to select regions and perform the permutations [109].

### Motif enrichment analysis

Transcription factor binding site enrichment within allele-specific ACRs (AS-ACRs) and SVs within ASE gene promoters were determined using MEME-ChIP with the parameters: (-meme-minw 5 -meme-maxw 30 -meme-nmotifs 10 -db ArabidopsisDAPv1.meme) [29]. Transcription factor binding motifs obtained from DNA affinity purification sequencing (DAP-seq) of *Arabidopsis thaliana* transcription factors [110] were used as the input database for motif identification. For AS-ACRs, the sequences for the maternal and paternal alleles were tested for motif-enrichment separately. For SVs in ASE gene promoters the sequence underlying the longer, consensus, ‘inserted’ allele was tested for motif-enrichment.

### Statistical inference of contribution of genomic features to allele-specific expression

An elastic-net regression model was developed to estimate the contribution of multiple factors to gene expression, including allele-specific expression. Overall gene expression was quantified using HTSeq-count with bias-corrected RNA-seq reads [111]. Genes that were outliers overall whole-genome sequencing read counts (1.5x interquartile range) were filtered prior to model fitting. For every gene, or those genes with significant ASE in both RNA-seq replicates, we gathered information for 56 factors (Additional File 1: Table S11). These factors include the overall read counts (or coverage) of each predictor as well as the log2 fold change of phased allele counts (Maternal / Paternal). For whole-genome sequencing reads, ChIP-seq reads, ATAC-seq reads and bisulfite-seq reads, the counts were classified as belonging to one of four regions surrounding each gene: genic (including exon and introns), promoter (< 1Kb from the transcription start site), upstream putative regulatory region (5 Kb upstream of promoter), and downstream putative regulatory region (5 Kb downstream of gene). Upstream and downstream regions were only considered if an ACR was detected in the window.

For the analysis of allelic differences in read coverage, the four regions of interest were set to NA if they did not contain at least one phased SNP. Additionally, regions were set to NA if their total SNP-based read counts were lower than the 10th percentile of SNP-based read counts for a given factor (ATAC-seq, ChIP-seq, BS-seq, etc). This filter was applied for each factor x region combination. Genes with a strong bias in whole-genome sequencing read counts (1.5x interquartile range) were filtered prior to model fitting (n=146). After filtering, the extent of missing data per factor ranged from 0 - 70% with a median of 11.3% missing data (Additional File 3: Tables S13 and S14). Elastic-net regression models are well suited to handle missing data and model fitting was performed with the glmnet package in R [112]. Model parameters were determined using 5-fold cross validation (Additional File 1: Table S12). For the model of overall expression, there was evidence of overfitting as 54 / 56 predictors were determined to be significant. To address this, we selected a shrinkage parameter (lambda) that reduces model complexity while maintaining a prediction error (mean-squared error) within one standard error of the model determined by 5 fold-cross validation (Additional File 1: Tables S15, S16; Additional File 2: Figure S6)

### Effects of SNPs and indels

SnpEff v.5.0e was used to annotate the impact of each SNP. The genome annotation (GFF) and SNP VCF files were used to build the SnpEff database. To evaluate the impact of potentially deleterious variants on ASE and compare their effects with the other classes of variants, SNPs classified as having a “HIGH” impact were merged with SNPs in ASE genes. (Additional File 3: Table S15)

## Acknowledgements

We thank Dr. Robert Kreuger from the USDA-ARS Citrus and Date Germplasm Repository for providing access to ‘Fairchild’ for sampling and James Burnette and Venkateswari Jaganatha Chetty for sampling of leaves from ‘Fairchild’. This research was supported by USDA-NIFA Award # 2020-70029-33202 to J.E.S. and D.K.S; National Science Foundation award to R.J.S. (IOS01856627) and to S.R.W. and J.E.S. (IOS1027542). J.E.S. is a CIFAR fellow in the Fungal Kingdom: Threats and Opportunities program. I.A.D is a fellow in the Plants-3D NSF National Research Traineeship Program (DBI-1922642).

## Availability of data and materials

Raw sequencing data used for genome assembly and annotation (Pacbio, 10x Genomic linked-reads, illumina whole-genome and RNA sequencing) are available as an NCBI BioProject (PRJNA357623) under the accession number SAMN06146124. The raw sequencing data used in the analyses of chromatin accessibility, histone modification, and DNA methylation are available as an NCBI BioProject (PRJNA1062830). Custom scripts used in analyses are available on GitHub [113].

## Additional file 2: Supplementary figure legends

**Figure S1.** Circos plot depicting the nine chromosomes of the ‘Fairchild’ genome assembly. **a)** Chromosomes. **b)** Genome-wide distribution of transposable elements. **c)** Genome-wide distribution of genes.

**Figure S2.** Chromatin landscape of genic accessible chromatin regions (ACRs). Metaplots were generated to summarize regional patterns of chromatin state, typically centered on the summit of an open chromatin region, which is defined as the 50 bp window with the highest tn5 integration frequency. For each data set, read coverage was scaled from 0 to 1 by dividing the read coverage at each site by the 98th quantile of sequencing coverage per sample [19]).

**Figure S3.** Chromatin landscape of proximal (within 2 Kb of nearest gene) accessible chromatin regions (ACRs). Metaplots were generated to summarize regional patterns of chromatin state, typically centered on the summit of an open chromatin region, which is defined as the 50 bp window with the highest tn5 integration frequency. For each data set, read coverage was scaled from 0 to 1 by dividing the read coverage at each site by the 98th quantile of sequencing coverage per sample. ACRs were clustered into five groups based on their patterns of histone modification.

**Figure S4.** Chromatin landscape of proximal (more than 2 Kb of nearest gene) accessible chromatin regions (ACRs). Metaplots were generated to summarize regional patterns of chromatin state, typically centered on the summit of an open chromatin region, which is defined as the 50 bp window with the highest tn5 integration frequency. For each data set, read coverage was scaled from 0 to 1 by dividing the read coverage at each site by the 98th quantile of sequencing coverage per sample. ACRs were clustered into five groups based on their patterns of histone modification.

**Figure S5.** Modeling of overall gene expression of all genes in the ‘Fairchild’ genome. Coefficients for significant predictors of overall expression (log2(TPM)) of all genes (n = 30,724) ordered by magnitude (R = 65.15). Factors are partitioned by genomic region and colored to indicate whether they reside in genes, promoters (1kb of TSS), or upstream/downstream putative regulatory regions of the focal gene (ACRs present 5 Kb upstream of promoter / 5 Kb downstream of gene). A summary of all models used is available in additional file 1: Table S12.

**Figure S6.** Determination of shrinkage parameter (*λ*) to minimize model complexity while maintaining predictive accuracy for overall expression (log2(TPM)) of all genes. The relationship between the shrinkage parameter (*λ*) and prediction accuracy (mean-squared error) during 5-fold cross validation with a constant tuning parameter (ɑ = 0.70853). The dotted vertical lines represent the range of *λ* values that maintain a prediction accuracy within one standard-error of the model chosen through the initial cross-validation. The red vertical line indicates the *λ* value chosen to minimize model complexity while maintaining performance. A summary of all models used is available in additional file 1: Table S12.

**Figure S7.** Modeling of overall gene expression of all genes in the ‘Fairchild’ genome after increasing shrinkage parameter (*λ*). Coefficients for significant predictors of overall expression (log2(TPM)) of all genes (n = 30,724) ordered by magnitude (R = 64.71). Factors are partitioned by genomic region and colored to indicate whether they reside in genes, promoters (1kb of TSS), or upstream/downstream putative regulatory regions of the focal gene (ACRs present 5 Kb upstream of promoter / 5 Kb downstream of gene). A summary of all models used is available in additional file 1: Table S12.

**Figure S8.** Distribution of size of deletions detected using 10x Genomics linked-reads (n = 2,463).

**Figure S9.** Promoters of ASE genes are enriched for deletions. The number of unique overlaps between promoters of ASE genes and deletions (green vertical line) was compared to 1,000 permutations of promoters from all genes in the Fairchild genome (p < 0.001). The black vertical line indicates the mean of the 1,000 permutations and the red vertical line indicates a significance threshold (p = 0.05).

**Figure S10.** Transposable element content of structural variation in gene promoters. **a)** The proportion of either: all promoters, all structural variants (deletions), all annotated transposable elements, or structural variants in promoters of ASE genes that belong to specific DNA transposon families or retrotransposons **(b)**.

**Figure S11.** AZF1 motif occurrence in structural variants within promoters of ASE genes. **a)** A motif enrichment analysis of sequence removed in the ‘deleted’ allele of 104 ASE genes with deletions in their promoter identified four significantly enriched motifs, with the most abundant (n=28) matching the binding site of zinc-finger protein 1 (AZF1), a zinc-finger protein that acts as a transcriptional repressor [25]. Comparison of allele-specific expression of the deletion containing allele ((Hap_Deletion_)^RNA^) of genes with the motif in the ‘deleted’ allele (n=28) compared to those without the motif (n = 78) indicates that there is not a significant effect of motif presence on allele expression (10,000 permutations, p = 0.27). **b)** Distribution of the difference in mean expression of the deletion containing allele ((Hap_Deletion_)^RNA^) for 10,000 permuted groups of genes selected from the 104 ASE genes with promoter deletions. The red dotted-line represents the observed difference in means between genes with the AZF1 motif in the deleted allele versus those without the motif.

**Figure S12.** The chromatin landscape of allele-specific ACRs is similar to all ACRs. Meta plots were generated to summarize regional patterns of chromatin accessibility and histone modification surrounding ACRs. ACRs were partitioned into three groups, all ACRs, GENEiASE ACRs (those with sufficient coverage and polymorphism for detection of ASE), and allele-specific ACRs (AS-ACRs). ACRs were scaled to a 1000 bp region, as well as their 2kb flanking regions using deepTools v.3.5.1 [104] and signal densities for each feature were plotted across these regions in 50 bp windows. For each data set, read coverage was scaled from 0 to 1 by dividing the read coverage at each site by the 98th quantile of sequencing coverage per sample [19].

**Figure S13.** The relationship between allele-accessibility of AS-ACRs and allele-expression of ASE genes. AS-ACRs were paired with the nearest ASE gene if they were present in the same phase-block (n=122, median distance = 7447.5 bp). Allele-accessibility of AS-ACRs represented as the ratio of maternal: paternal ATAC-seq reads is positively correlated with allele-specific expression of neighboring genes (R = 0.46, p = 1.9e-07). Points are colored by the distance between the AS-ACRs and its nearest ASE gene.

**Figure S14.** AS-ACRs are enriched for DNA transposons. **a)** The number of unique overlaps between AS-ACRs and DNA transposons (green vertical line) was compared to 1,000 permutations from all ACRs in the Fairchild genome (p = 0.007). The black vertical line indicates the mean of the 1,000 permutations and the red vertical line indicates a significance threshold (p = 0.05). **b)** The proportion of AS-ACRs overlapping transposons belonging to specific families.

**Figure S15.** AS-ACRs are not enriched for retrotransposons. **a)** The number of unique overlaps between AS-ACRs and retrotransposons (green vertical line) was compared to 1,000 permutations from all ACRs in the Fairchild genome (p = 0.5345). The black vertical line indicates the mean of the 1,000 permutations and the red vertical line indicates a significance threshold (p = 0.05).

**Figure S16.** The relationship between allele-accessibility of AS-ACRs and allele-expression of ASE genes decays over longer distances. **a)** ASE genes were paired with their nearest AS-ACR and only ACR-gene pairs within the same phase block were retained. ACR-gene pairs were then split into 25 quantiles based on the distance between gene and ACR. For each quantile, the spearman correlation coefficient (rho) of allele-accessibility and allele-expression of gene-ACR pairs (n=30) was calculated. As a control, ASE genes were paired with their nearest ACR, regardless of whether the ACR has significant allele-specificity. The spearman correlation coefficient was calculated for control ACR-gene pairs (n=30), and we reported the average of 1000 permutations of ACR-gene pairs. The vertical lines indicate the 95% confidence interval for the spearman correlation coefficient (rho).

**Figure S17.** Interspersed repeat landscape. The interspersed repeat landscape, revealing the distribution and divergence of repeat elements across the genome. The percentages of the genome composed of various repeat elements (Y-axis) are categorized according to their Kimura substitution level (X-axis; CpG adjusted values ranging from 0 to 50), each bar in the plot represents a 1% sequence divergence bin, adjusted for CpG Kimura divergence. The landscape is represented by a stacked bar plot, where each color corresponds to a specific class of repeats. The plot illustrates the copy-divergence analysis of these elements, with older copies indicated by higher Kimura values towards the right, and more recent copies towards the left, indicating the evolutionary dynamics of the repetitive elements within the genome.

