## Additional File 2 Supplemental Figures for "Haplotype phased genome of ‘Fairchild’ mandarin highlights influence of local chromatin state on gene expression"

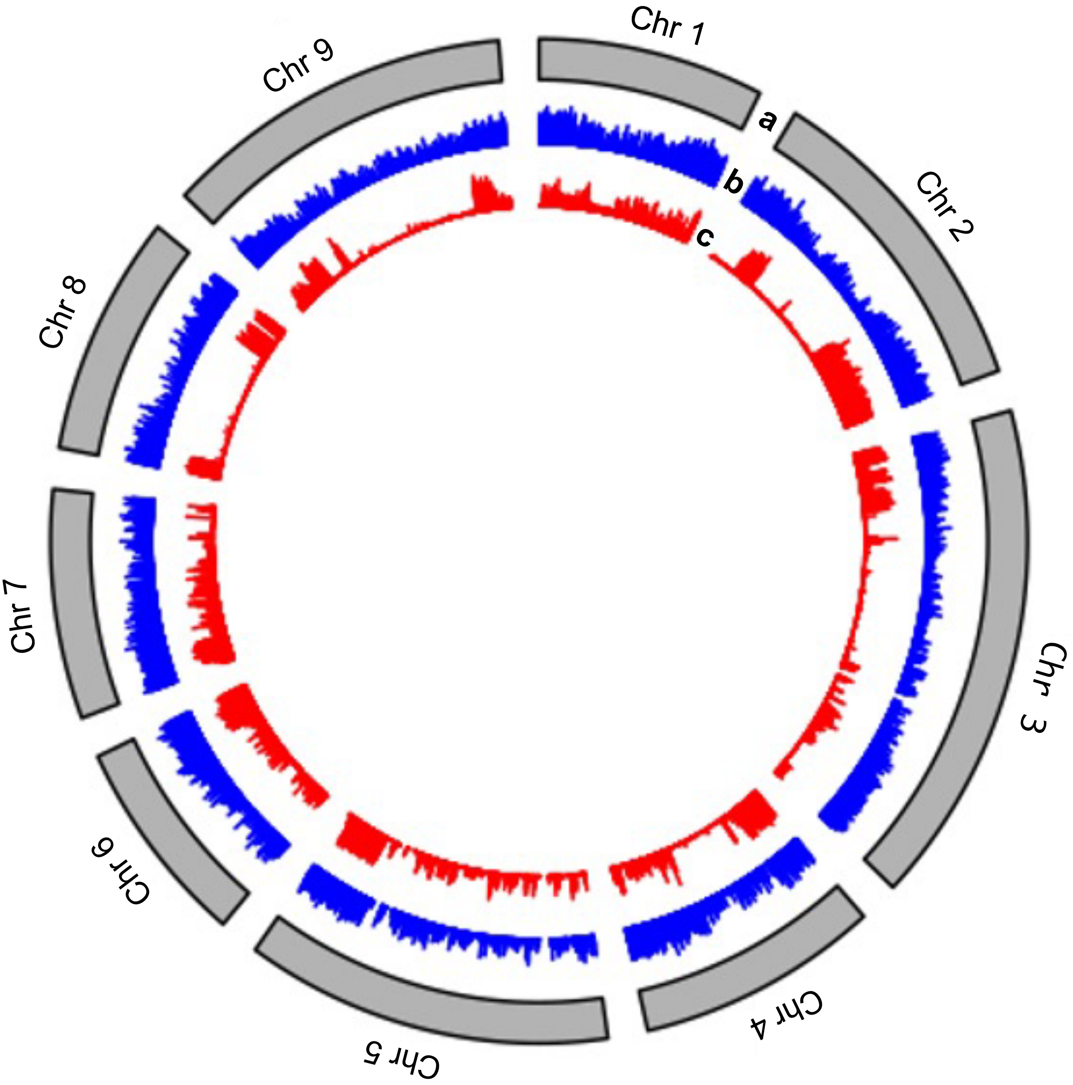
Chr 1
Chr 2
Chr 3
Chr 4
Chr 5
Chr 6
Chr 7
Chr 8
Chr 9
a
b
c

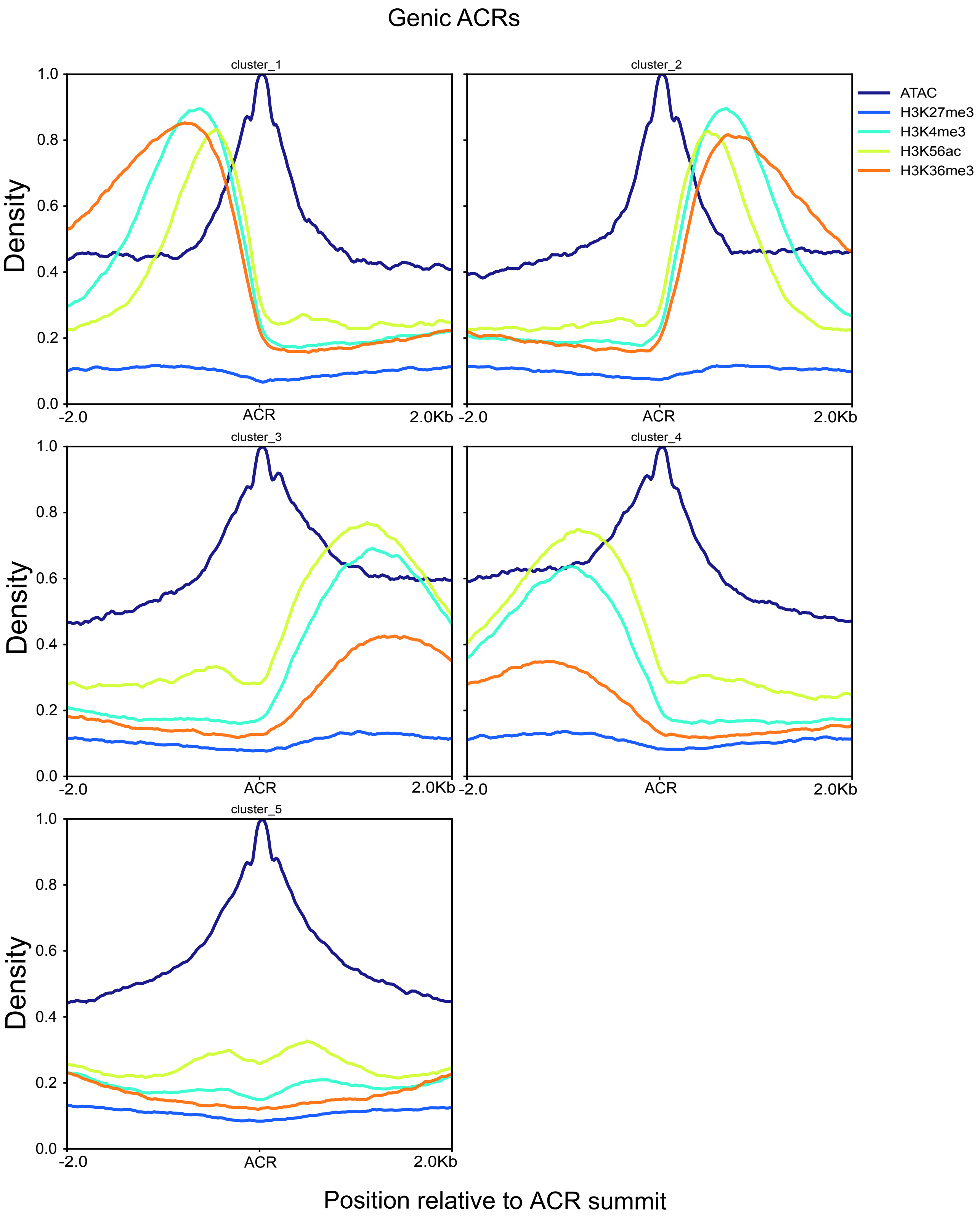

Genic ACRs
cluster_1
cluster_2
cluster_3
cluster_4
cluster_5
ATAC
H3K27me3
H3K4me3
H3K56ac
H3K36me3
Density
0.0
0.2
0.4
0.6
0.8
1.0
-2.0
ACR
2.0Kb
Position relative to ACR summit

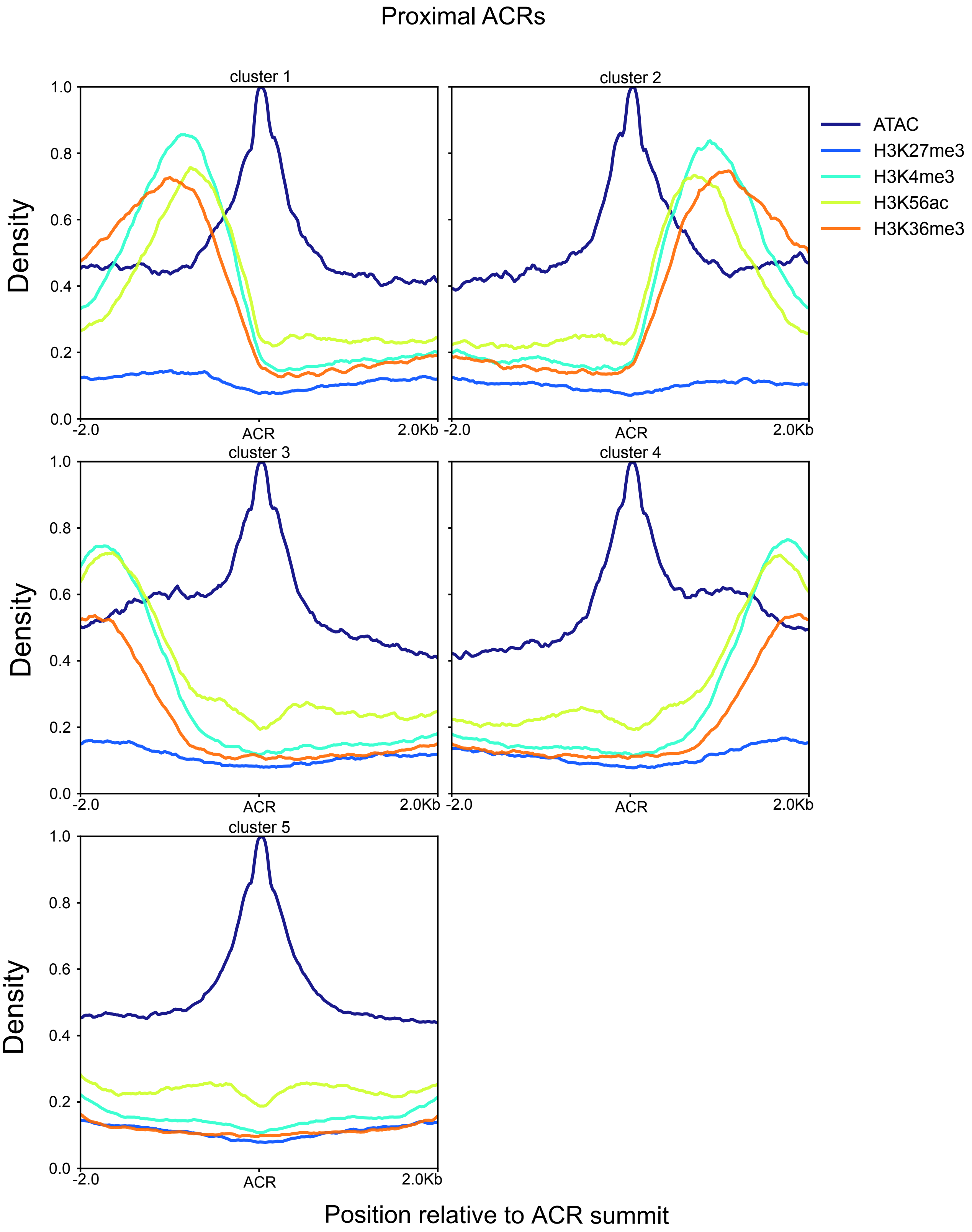

Proximal ACRs
cluster 1
cluster 2
cluster 3
cluster 4
cluster 5
ATAC
H3K27me3
H3K4me3
H3K56ac
H3K36me3
Density
1.0
0.8
0.6
0.4
0.2
0.0
-2.0
ACR
2.0Kb
Position relative to ACR summit

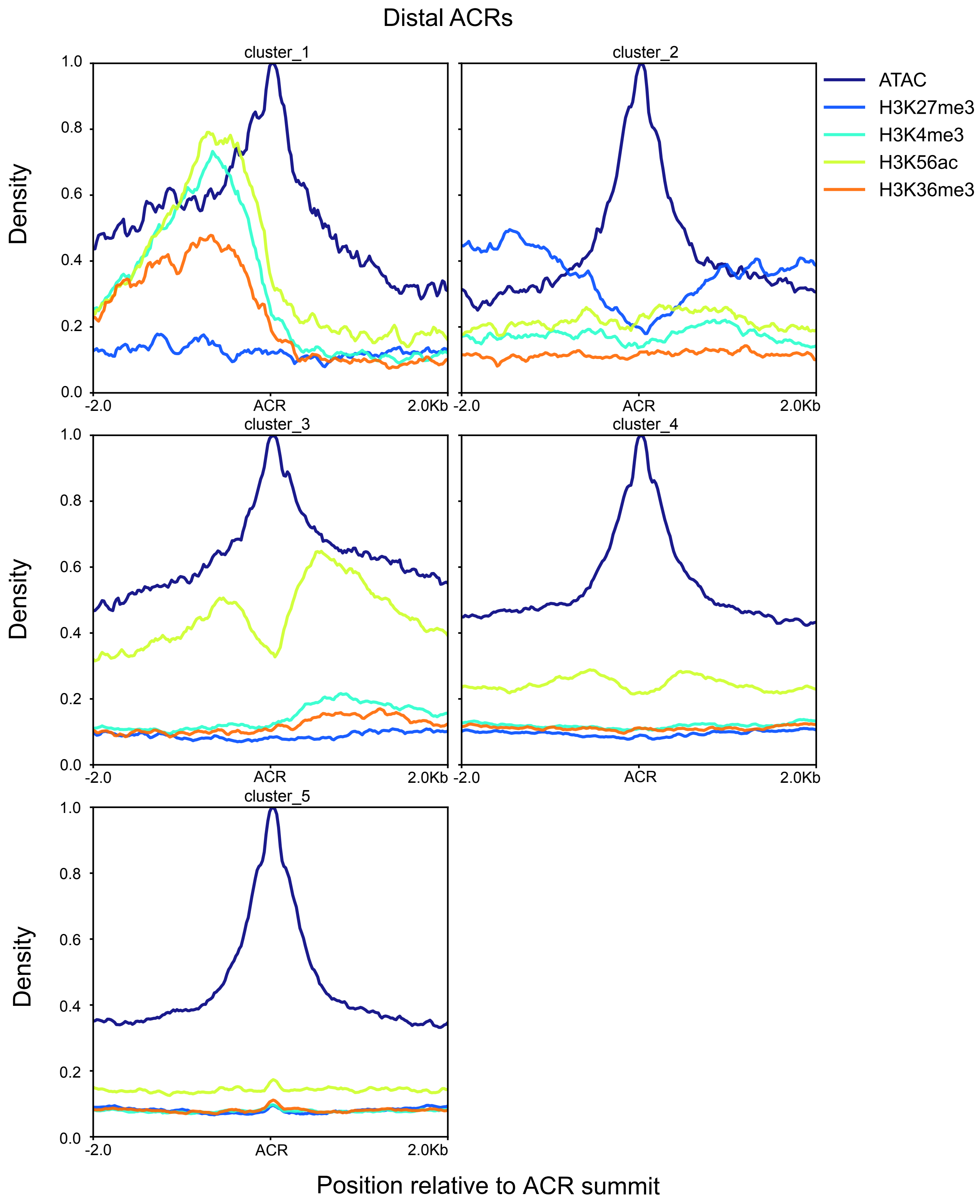
Distal ACRs
cluster_1
cluster_2
cluster_3
cluster_4
cluster_5
ATAC
H3K27me3
H3K4me3
H3K56ac
H3K36me3
Density
1.0
0.8
0.6
0.4
0.2
0.0
-2.0
ACR
2.0Kb
Position relative to ACR summit

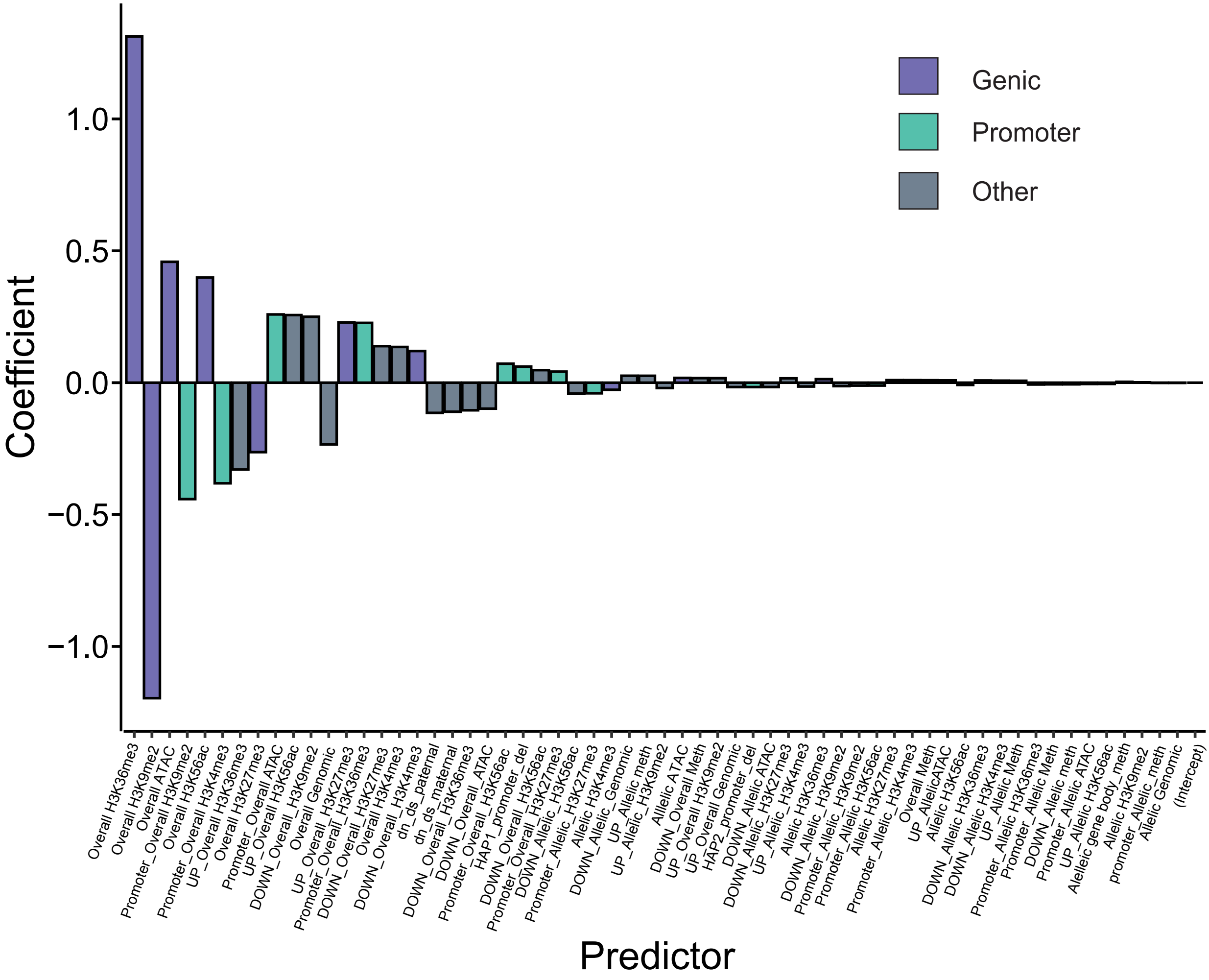

Genic
Promoter
Other
Coefficient
1.0
0.5
0.0
−0.5
−1.0
Predictor
Overall H3K36me3
Overall H3K9me2
Overall ATAC
Promoter_Overall H3K9me2
Overall H3K56ac
Promoter_Overall H3K4me3
UP_Overall H3K36me3
Overall H3K27me3
Promoter_Overall ATAC
UP_Overall H3K56ac
DOWN_Overall_H3K9me2
Overall Genomic
UP_Overall_H3K27me3
Promoter_Overall_H3K36me3
DOWN_Overall_H3K27me3
Overall H3K4me3
DOWN_Overall_H3K4me3
dn_ds_paternal
dn_ds_maternal
DOWN_Overall_H3K36me3
DOWN_Overall_ATAC
Promoter_Overall_H3K56ac
HAP1_promoter_del
DOWN_Overall_H3K56ac
Promoter_Overall_H3K27me3
DOWN_Allelic_H3K56ac
Promoter_Allelic_H3K27me3
Allelic H3K4me3
DOWN_Allelic_Genomic
UP_Allelic meth
UP_Allelic_H3K9me2
Alllelic ATAC
DOWN_Overall Meth
UP_Overall H3K9me2
UP_Overall Genomic
HAP2_promoter_del
DOWN_Allelic ATAC
DOWN_Allelic_H3K27me3
UP_Allelic_H3K4me3
Allelic H3K36me3
DOWN_Allelic_H3K9me2
Promoter_Allelic_H3K9me2
Promoter_Allelic H3K56ac
Allelic H3K27me3
Promoter_Allelic_H3K4me3
Overall Meth
UP_AllelicATAC
Allelic H3K56ac
DOWN_Allelic H3K36me3
DOWN_Allelic H3K4me3
UP_Allelic Meth
Promoter_Allelic H3K36me3
Promoter_Allelic Meth
DOWN_Allelic meth
Promoter_Allelic ATAC
UP_Allelic H3K56ac
Aleleic gene body_meth
Allelic H3K9me2
promoter_Allelic_meth
Allelic Genomic
(Intercept)

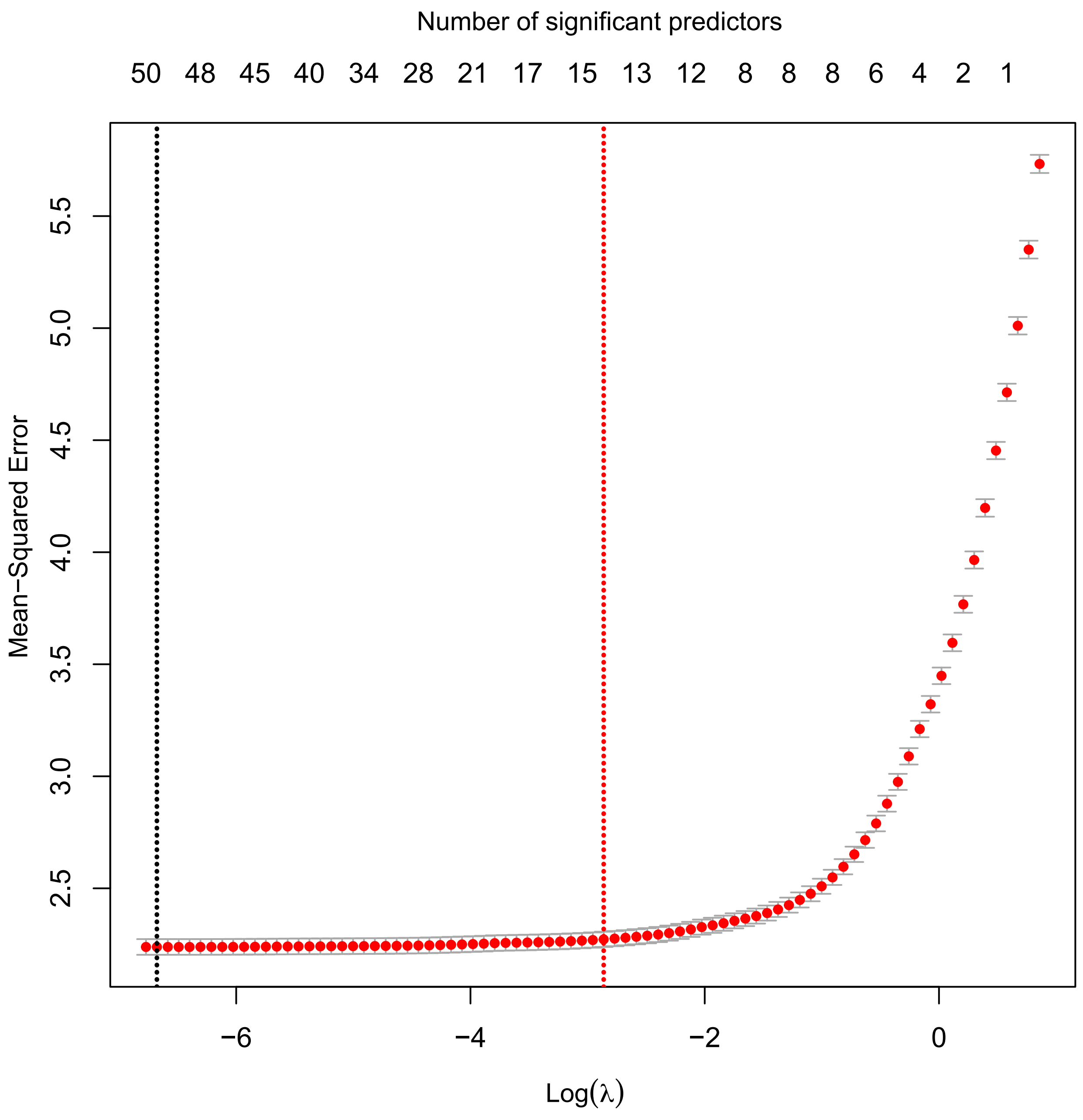

Number of significant predictors
50 48 45 40 34 28 21 17 15 13 12 8 8 8 6 4 2 1
Mean−Squared Error
2.5
3.0
3.5
4.0
4.5
5.0
5.5
−6 −4 −2 0
Log(λ)

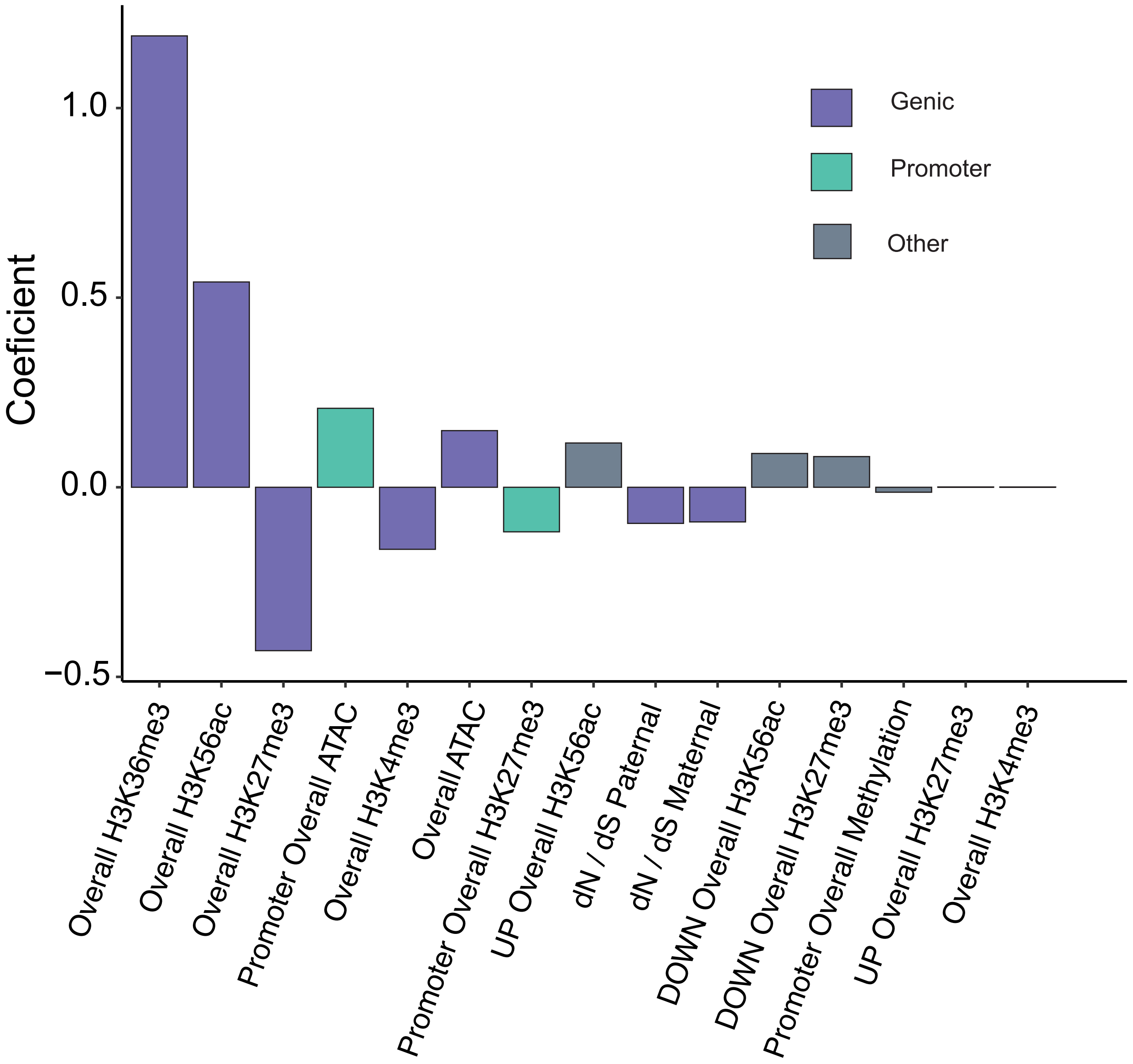
Coeficient
1.0
0.5
0.0
−0.5
Genic
Promoter
Other
Overall H3K36me3
Overall H3K56ac
Overall H3K27me3
Promoter Overall ATAC
Overall H3K4me3
Overall ATAC
Promoter Overall H3K27me3
UP Overall H3K56ac
dN / dS Paternal
dN / dS Maternal
DOWN Overall H3K56ac
DOWN Overall H3K27me3
Promoter Overall Methylation
UP Overall H3K27me3
Overall H3K4me3

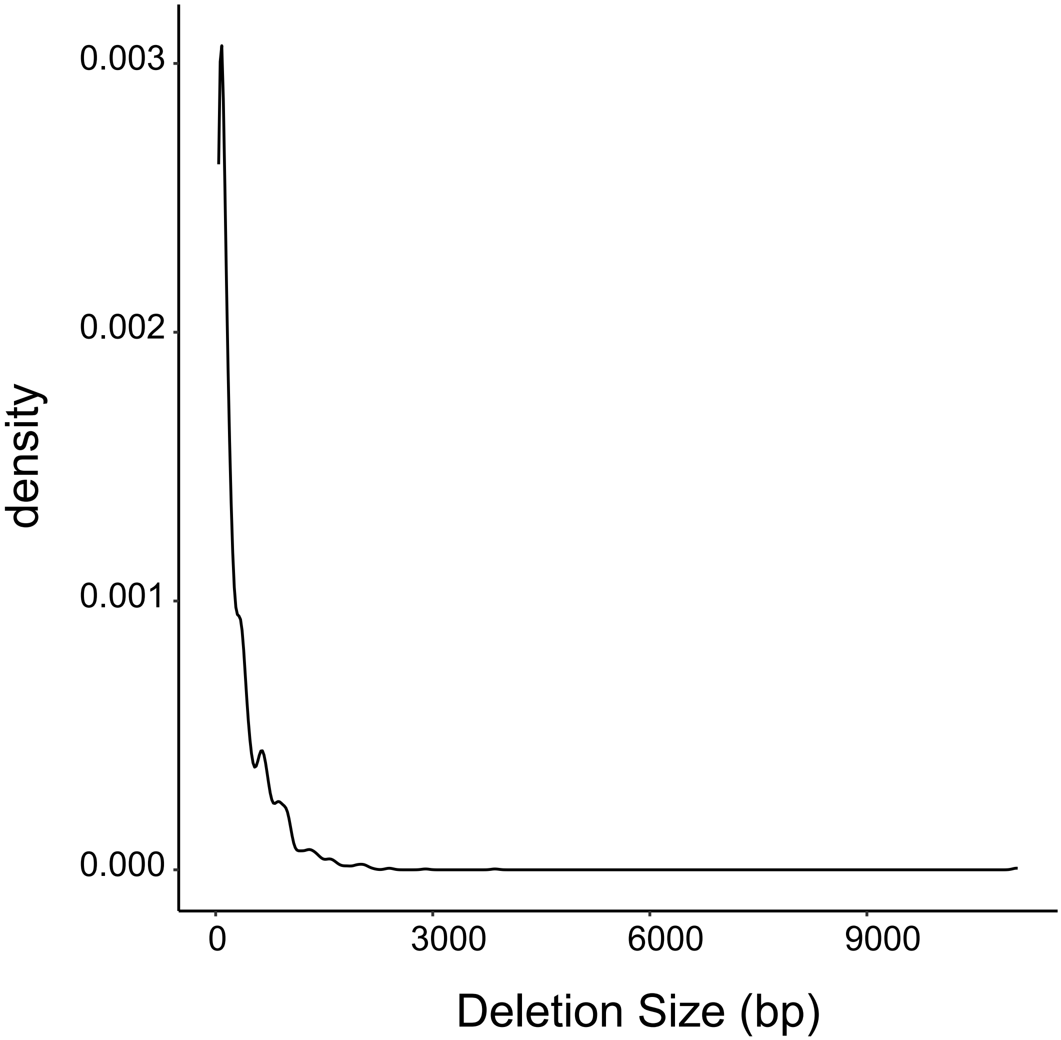
density
0.003
0.002
0.001
0.000
0
3000
6000
9000
Deletion Size (bp)

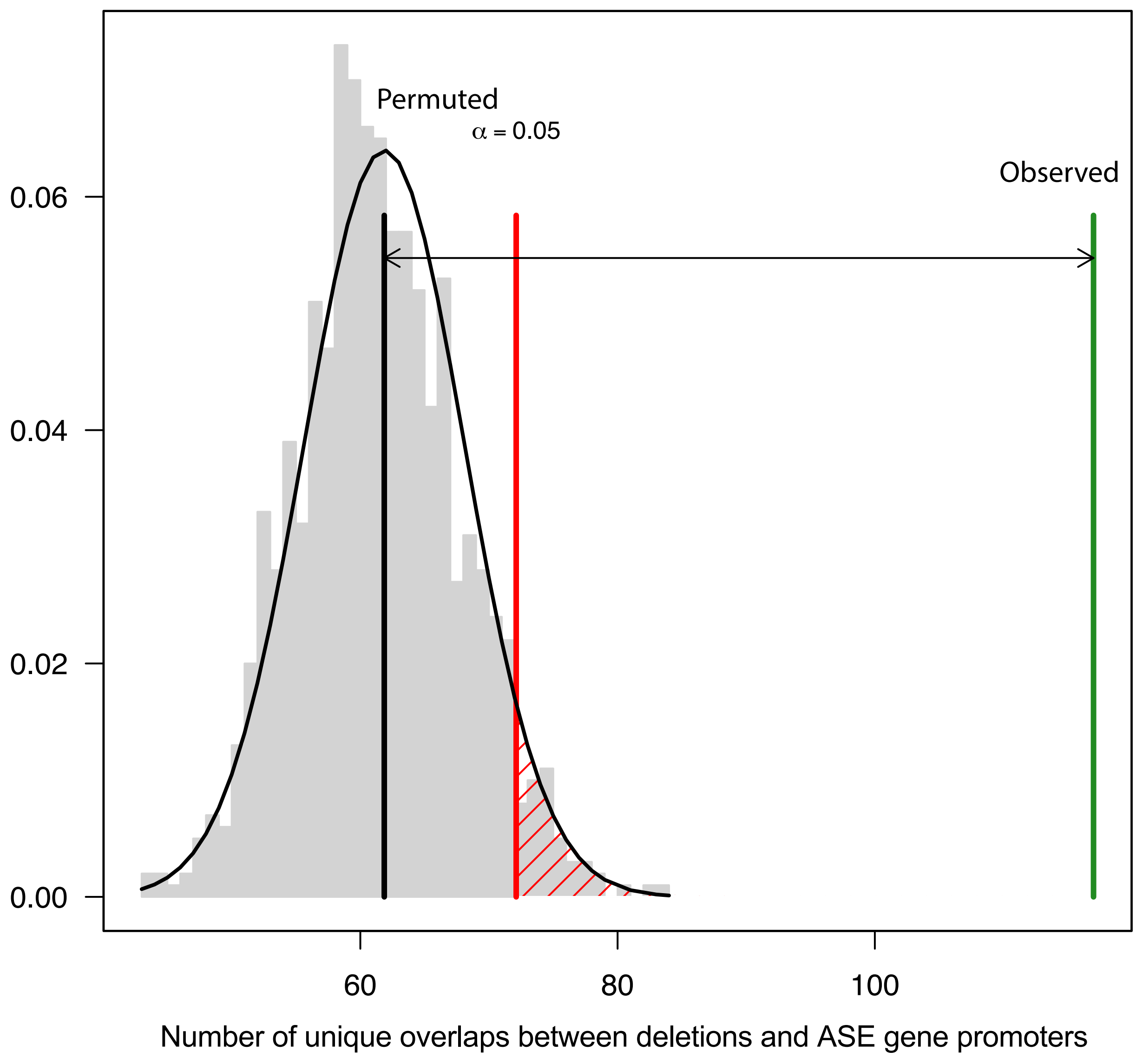

Permuted
α = 0.05
Observed
0.06
0.04
0.02
0.00
60
80
100
Number of unique overlaps between deletions and ASE gene promoters

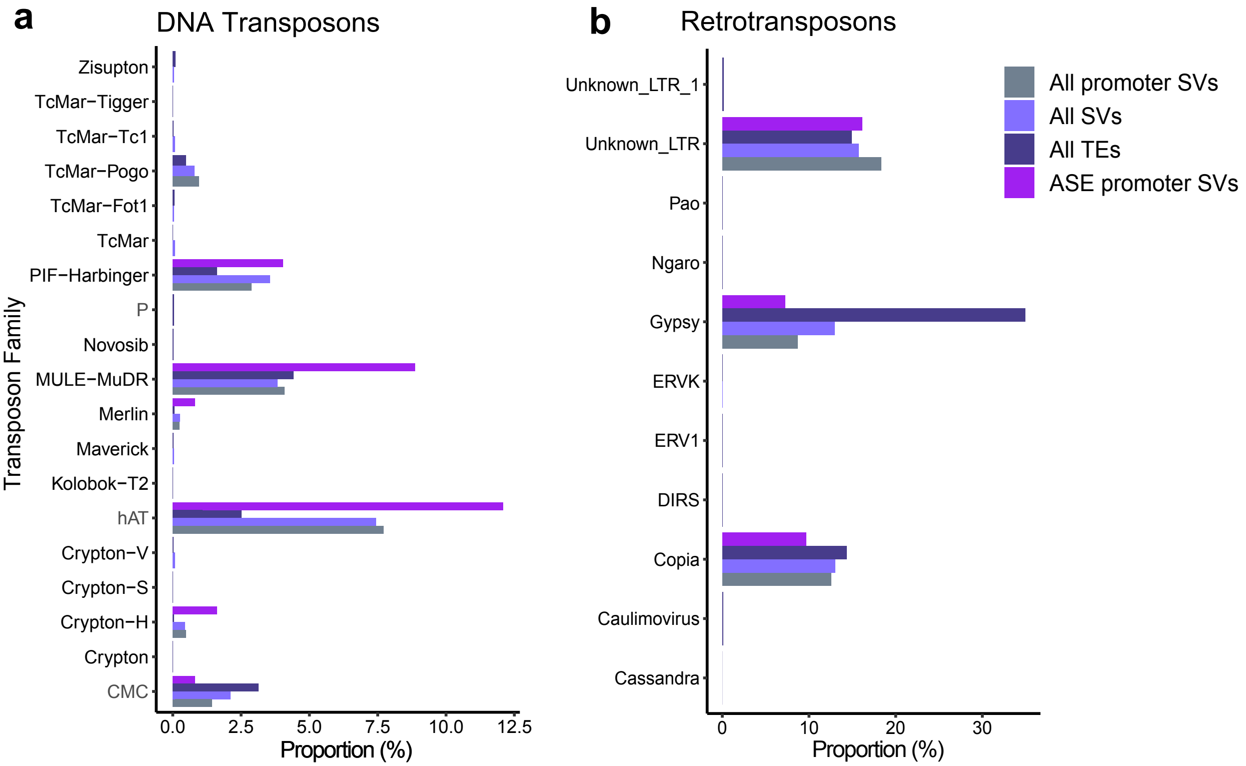
a
DNA Transposons
Transposon Family
Zisupton
TcMar−Tigger
TcMar−Tc1
TcMar−Pogo
TcMar−Fot1
TcMar
PIF−Harbinger
P
Novosib
MULE−MuDR
Merlin
Maverick
Kolobok−T2
hAT
Crypton−V
Crypton−S
Crypton−H
Crypton
CMC
0.0
2.5
5.0
7.5
10.0
12.5
Proportion (%)
b
Retrotransposons
Unknown_LTR_1
Unknown_LTR
Pao
Ngaro
Gypsy
ERVK
ERV1
DIRS
Copia
Caulimovirus
Cassandra
0
10
20
30
Proportion (%)
All promoter SVs
All SVs
All TEs
ASE promoter SVs

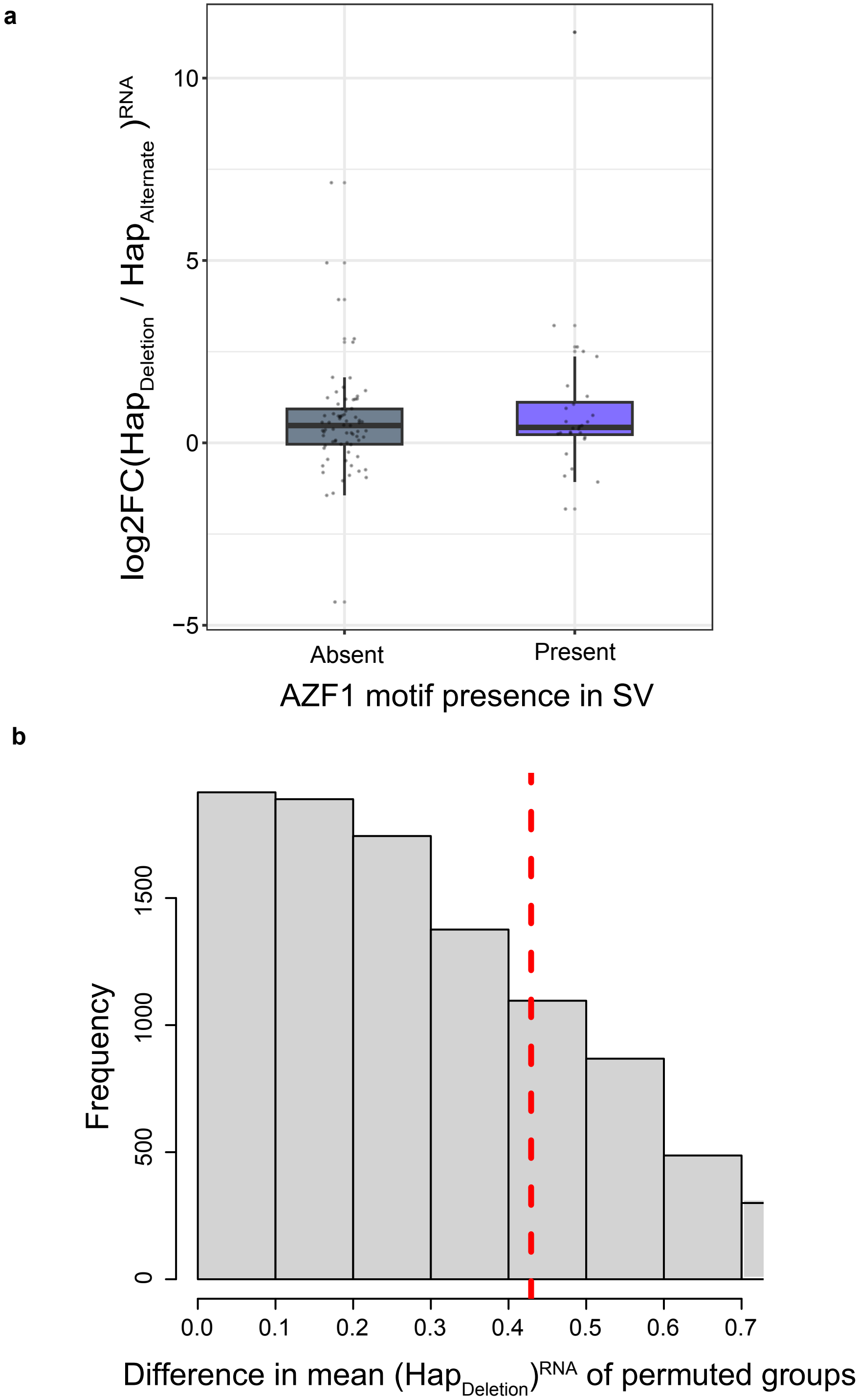
a
log2FC(Hap$_{Deletion}$ / Hap$_{Alternate}$)$^{RNA}$
10
5
0
−5
Absent
Present
AZF1 motif presence in SV
b
Frequency
0
500
1000
1500
0.0
0.1
0.2
0.3
0.4
0.5
0.6
0.7
Difference in mean (Hap$_{Deletion}$)$^{RNA}$ of permuted groups

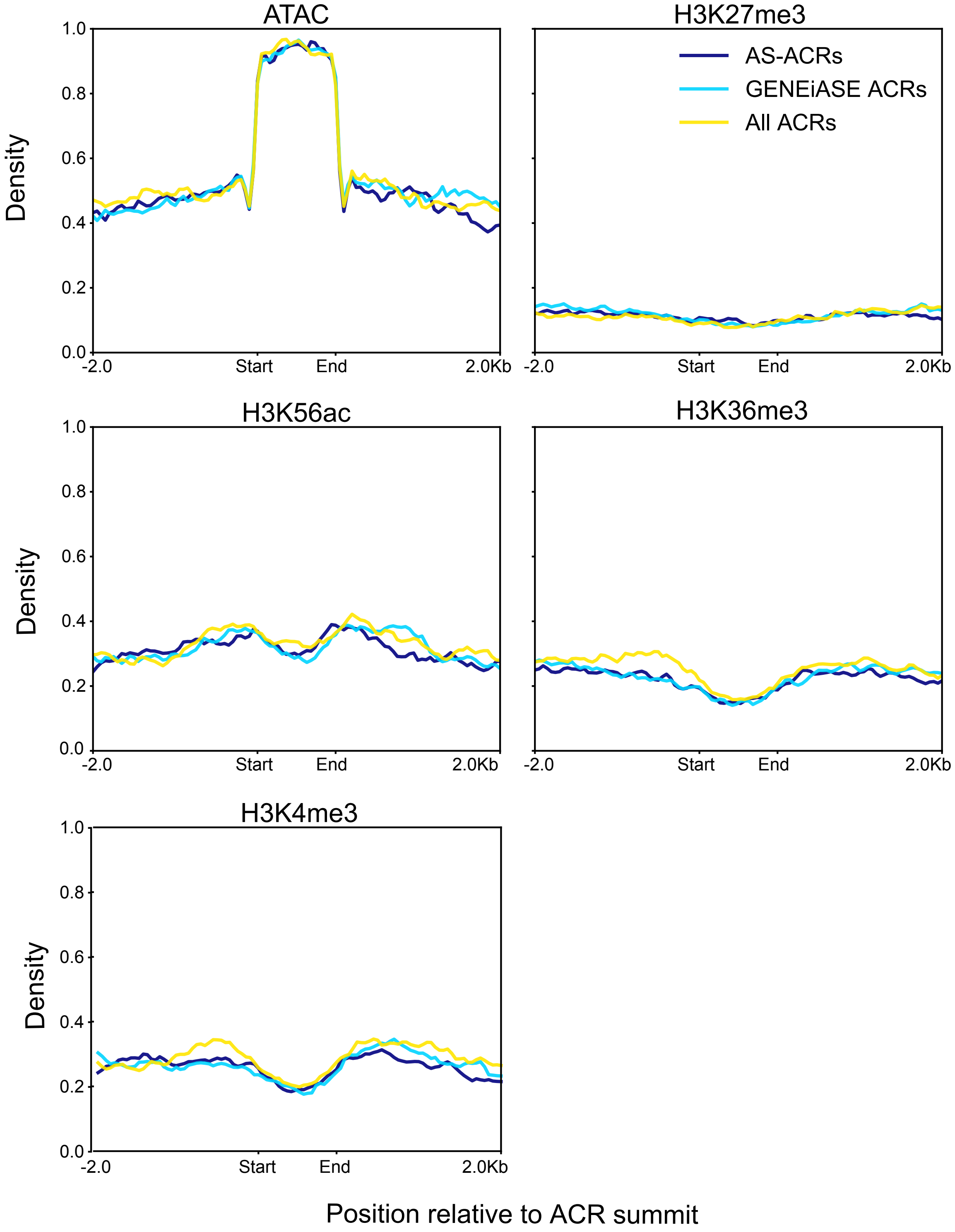
ATAC
H3K27me3
AS-ACRs
GENEiASE ACRs
All ACRs
H3K56ac
H3K36me3
H3K4me3
Density
1.0
0.8
0.6
0.4
0.2
0.0
-2.0
Start
End
2.0Kb
Position relative to ACR summit

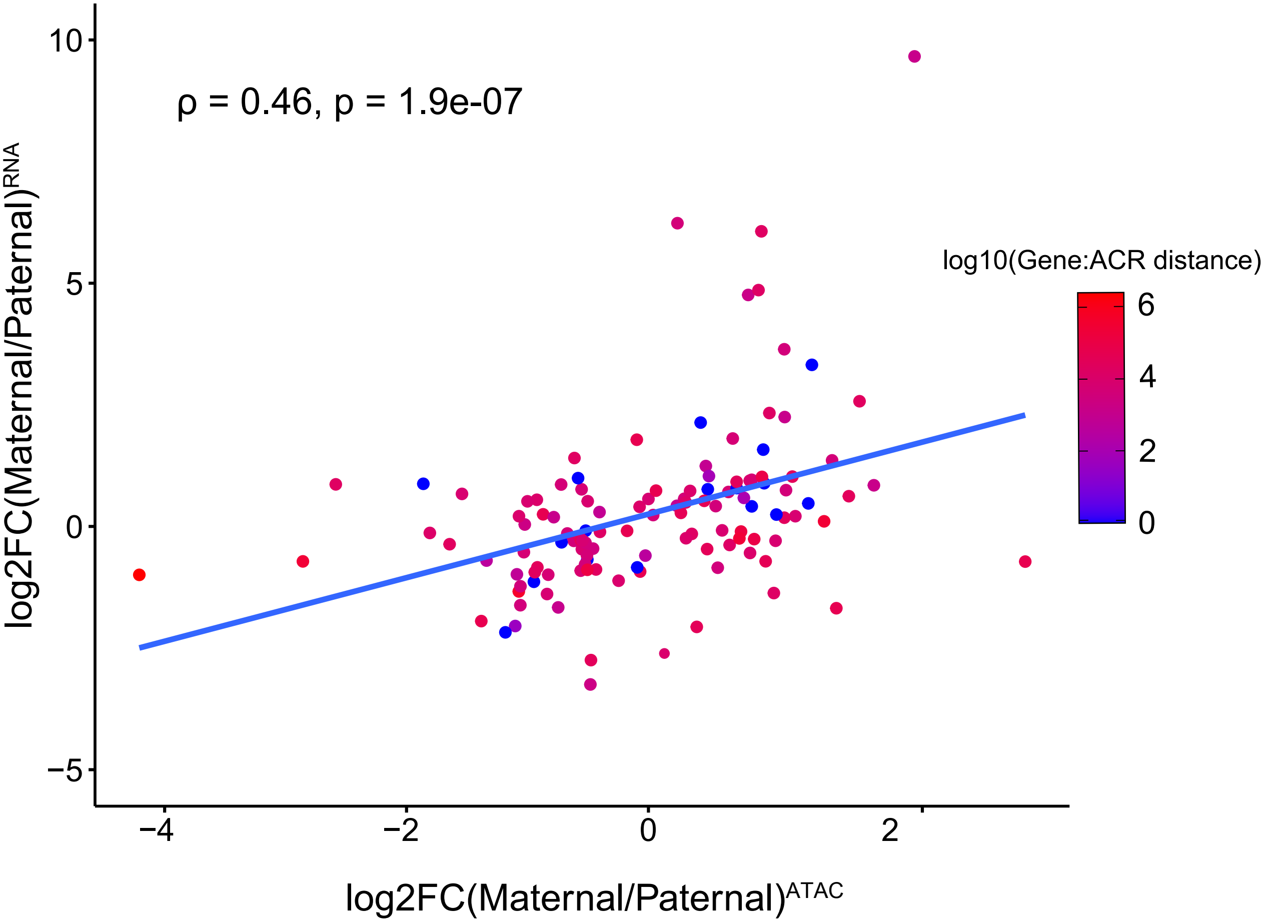
ρ = 0.46, p = 1.9e-07
log2FC(Maternal/Paternal)RNA
log2FC(Maternal/Paternal)ATAC
log10(Gene:ACR distance)
10
5
0
−5
−4
−2
0
2
6
4
2
0

a

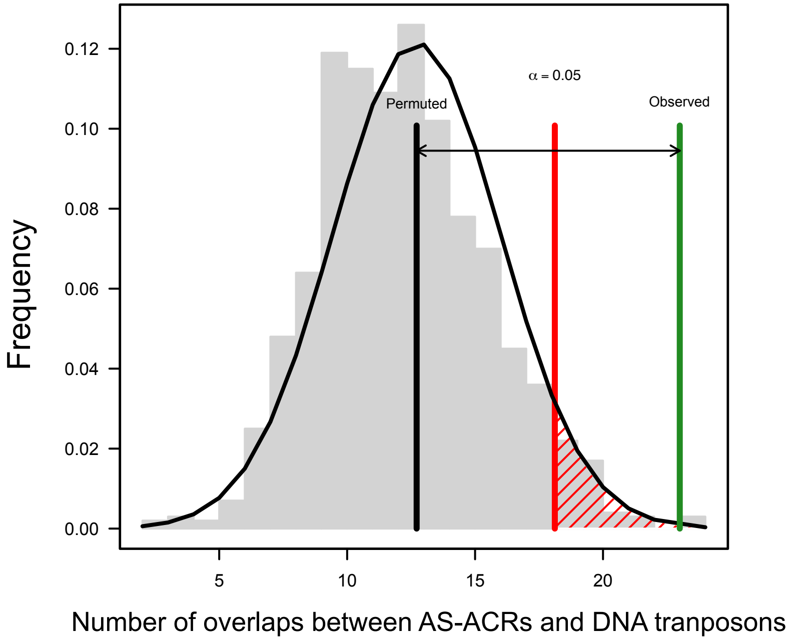

Frequency
0.12
0.10
0.08
0.06
0.04
0.02
0.00
5
10
15
20
Permuted
α = 0.05
Observed
Number of overlaps between AS-ACRs and DNA tranposons


b

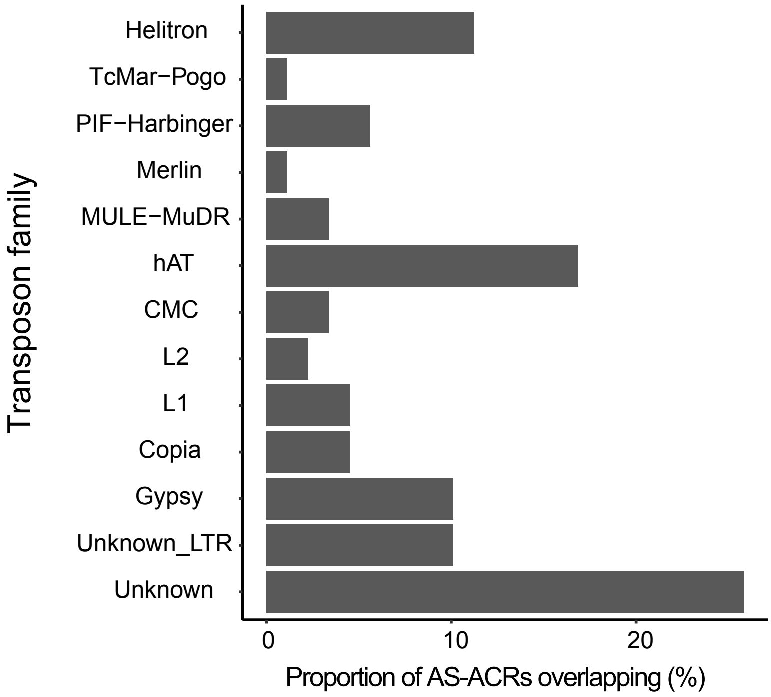

Transposon family
Helitron
TcMar−Pogo
PIF−Harbinger
Merlin
MULE−MuDR
hAT
CMC
L2
L1
Copia
Gypsy
Unknown_LTR
Unknown
0
10
20
Proportion of AS-ACRs overlapping (%)

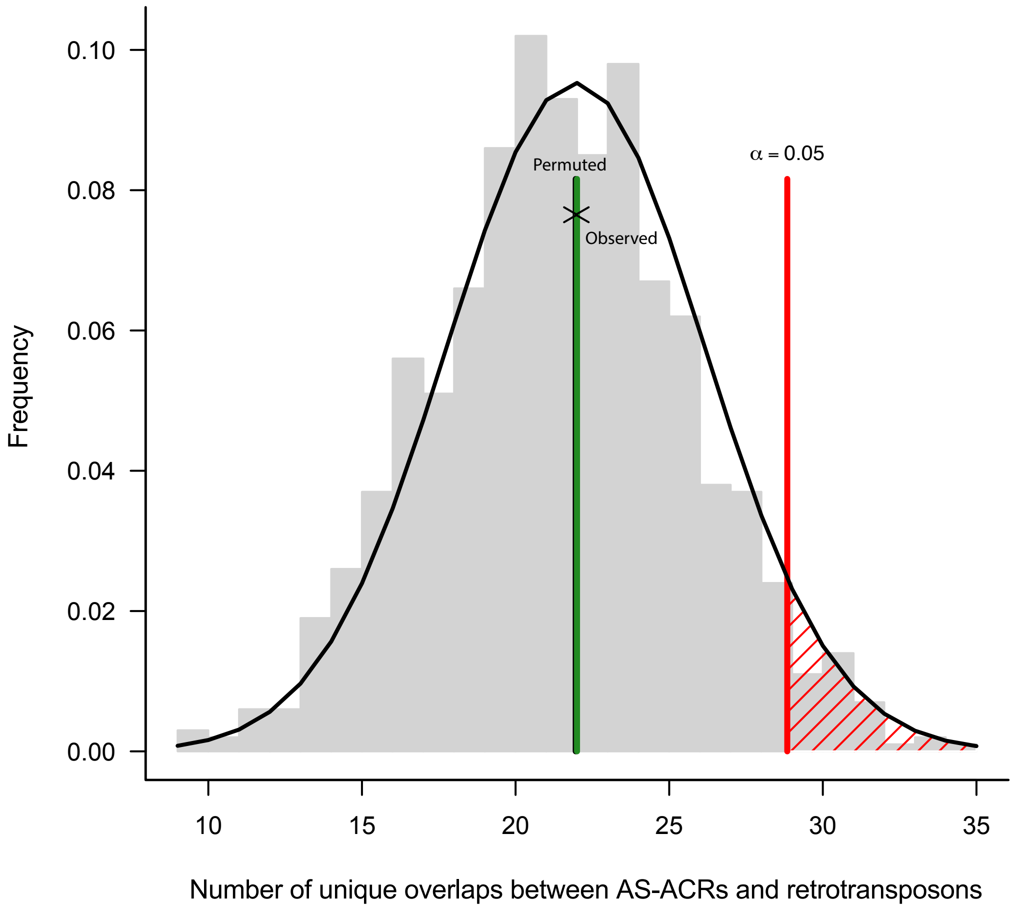

Frequency
0.10
0.08
0.06
0.04
0.02
0.00
Permuted
Observed
α = 0.05
10
15
20
25
30
35
Number of unique overlaps between AS-ACRs and retrotransposons

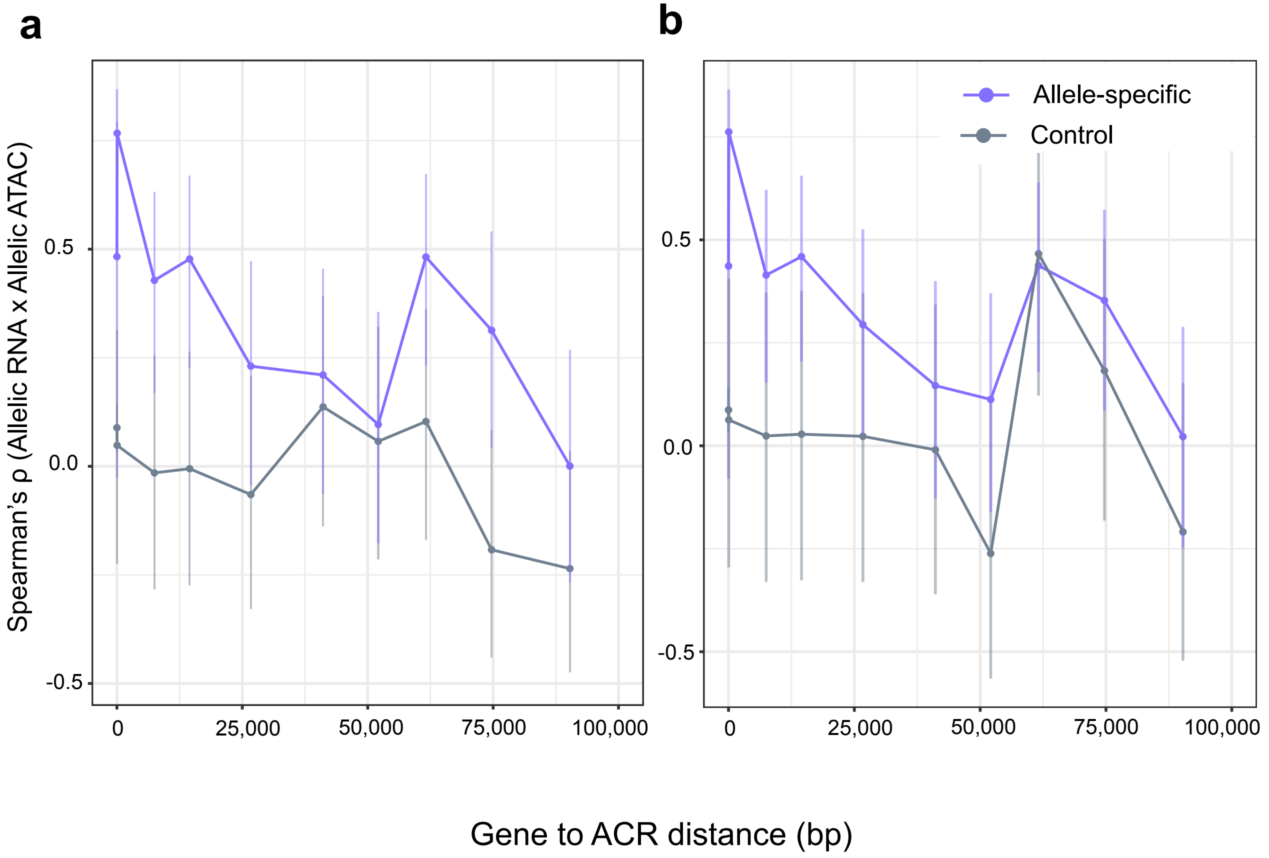
a
b
Spearman's ρ (Allelic RNA x Allelic ATAC)
0.5
0.0
-0.5
0
25,000
50,000
75,000
100,000
Allele-specific
Control
Gene to ACR distance (bp)

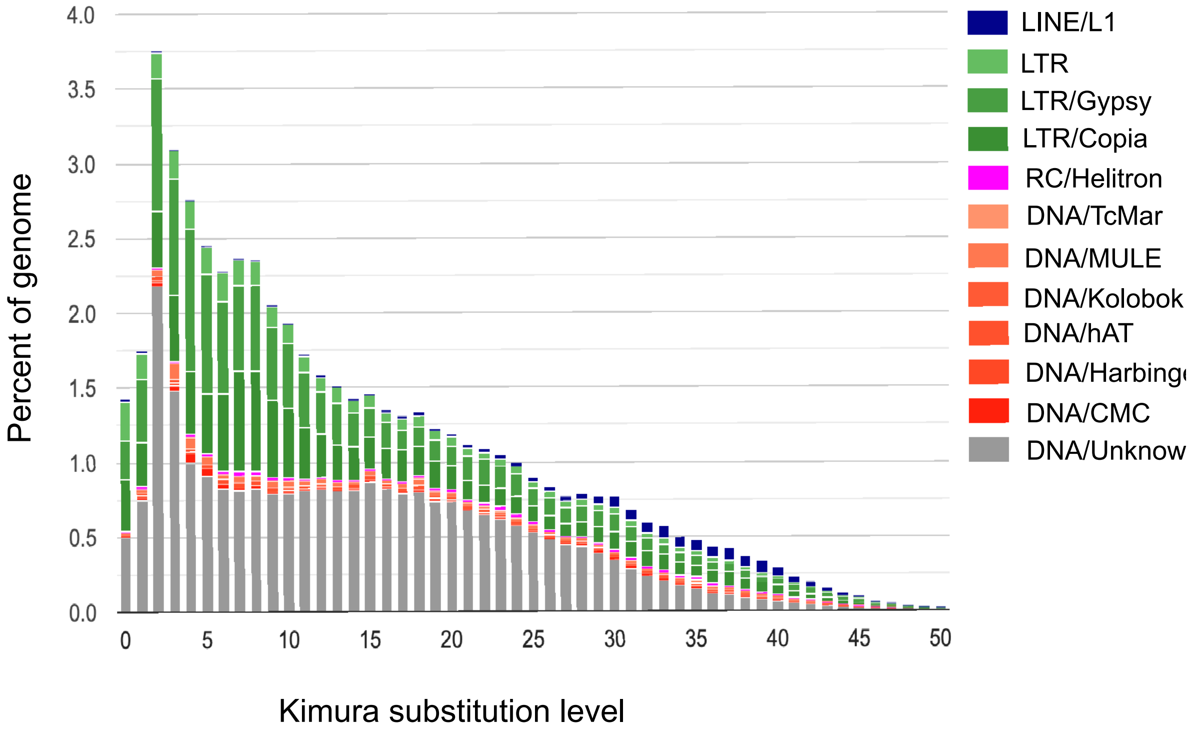

4.0
3.5
3.0
2.5
2.0
1.5
1.0
0.5
0.0
Percent of genome
0
5
10
15
20
25
30
35
40
45
50
Kimura substitution level
LINE/L1
LTR
LTR/Gypsy
LTR/Copia
RC/Helitron
DNA/TcMar
DNA/MULE
DNA/Kolobok
DNA/hAT
DNA/Harbing
DNA/CMC
DNA/Unknow
